# *Vibrio parahaemolyticus* quorum sensing controls phage VP882 transmission

**DOI:** 10.64898/2026.03.16.712183

**Authors:** Molly R. Sargen, Bonnie L. Bassler

## Abstract

Quorum sensing is a cell-to-cell communication process bacteria use to orchestrate collective behaviors. Quorum sensing involves the production, release, and detection of extracellular signal molecules called autoinducers. Some temperate phages can monitor bacterial autoinducers enabling them to track the abundance of potential host cells in the vicinity. Quorum-sensing-responsive phages can preferentially launch the transition from lysogeny to lytic replication at high cell density, presumably maximizing transmission. Once the phage lytic program is enacted, if nearby host cells are already lysogens, infections initiated by released virions could be nonproductive due to homoimmunity or superinfection exclusion mechanisms, posing a conundrum for temperate phages, including those that surveil quorum-sensing autoinducers. Here, we define host and phage components that influence transmission of the first discovered quorum-sensing-responsive phage, phage VP882, in populations of its host, *Vibrio parahaemolyticus*. We show that phage VP882 uses the K-antigen of serotype O3:K6 as its receptor. We demonstrate that host cells can prevent phage access to the O3:K6 K-antigen via quorum-sensing-control of the export of polysaccharides that shield the K-antigen from the phage at high cell density. We discover that phage VP882 can superinfect and superlysogenize *V. parahaemolyticus*, overcoming the challenge of detecting whether or not potential hosts are lysogens. Following superlysogenization, recombination of the resident and newly infecting phage genomes can occur possibly promoting phage VP882 genome diversification.

**Importance:** A longstanding mystery is how temperate phages optimally time the launch of their lytic cascades to maximize spread. Quorum-sensing-responsive phages can preferentially execute their lytic replication programs at high host cell density, which in principle should foster transmission. However, if nearby host cells are already lysogens, infections initiated by released virions could be nonproductive due to homoimmunity or superinfection exclusion. We define host and phage components influencing transmission of the quorum-sensing-responsive phage VP882 in *Vibrio parahaemolyticus* populations. Phage VP882 uses the O3:K6 K-antigen as its receptor. Host cells prevent phage infection via quorum-sensing-controlled export of polysaccharides that shield the K-antigen at high cell density. We discover that phage VP882 can superinfect and superlysogenize *V. parahaemolyticus*, overcoming the challenge of detecting whether or not potential hosts are lysogens. These findings reveal how phages can capitalize on interception of host quorum-sensing cues to maximize their reproductive success.

## Introduction

Temperate phages alternate between lytic and lysogenic lifestyles^1^. When a phage is lytic, it replicates, kills its current host cell, and spreads to new cells. During lysogeny, the phage establishes a stable relationship with its host and is transmitted vertically as a prophage. Transitioning between lytic and lysogenic lifestyles enhances phage reproductive success by maximizing the potential for transmission^2^. Theory and experimental evidence suggest that phage transmission through the lytic route is often favored when susceptible hosts are abundant while lysogeny is preferred when potential new hosts are rare^3,4^. Quorum sensing (QS) is a cell-to-cell communication process in which bacteria produce, release, and detect extracellular signal molecules called autoinducers. QS enables bacteria to orchestrate collective behaviors^5^. Some temperate phages can monitor host QS autoinducers, allowing them to estimate host cell abundance and launch the lytic cycle at high cell density when many host cells are present^6,7^.

A conundrum all temperate phages face when launching the lytic cycle is whether host cells in the vicinal community are already infected or not. Encountering bacteria that are already lysogens is often a dead-end for a phage because the lytic repressor produced from the genome of the resident lysogenic phage suppresses transcription from the newly infecting phage^8,9^. This process is known as homoimmunity and is protective for both the bacterial host and the resident prophage^10^. In other cases, prophages encode superinfection exclusion mechanisms that prevent infection of already lysogenized cells^11,12^. Superinfection immunity often occurs through alteration of the expression or function of the phage receptor on the host cell surface^13–16^. Superinfection exclusion protects lysogenic host cells and, unlike in the case of homoimmunity, also preserves the population of free phage virions so they are available to infect naïve hosts^17^.

Homoimmunity and superinfection exclusion mechanisms potentially threaten the reproductive success of QS-responsive prophages that launch their lytic cycles at high cell densities.

Phage VP882, the first phage discovered to surveil a host QS autoinducer, infects the pathogen *Vibrio parahaemolyticus*. *V. parahaemolyticus* has two QS systems (Figure 1). First, the VqmA-VqmR QS pathway is controlled by the DPO autoinducer^18^. At high cell density, the VqmA transcription factor binds DPO, and the DPO-VqmA complex activates expression of *vqmR* encoding the VqmR small RNA (sRNA). VqmR promotes expression of genes required for group behaviors. Second, the LuxO-driven QS cascade is responsive to three autoinducers that are detected by three membrane bound receptors and, via a phosphorylation-dephosphorylation cascade, the receptors modulate the activity of the LuxO transcription factor^5^. At low cell density, LuxO is phosphorylated and it controls a regulon underpinning individual behaviors. At high cell density, LuxO is dephosphorylated, and under this condition, the regulon responsible for group traits is enacted.

**Figure 1:**
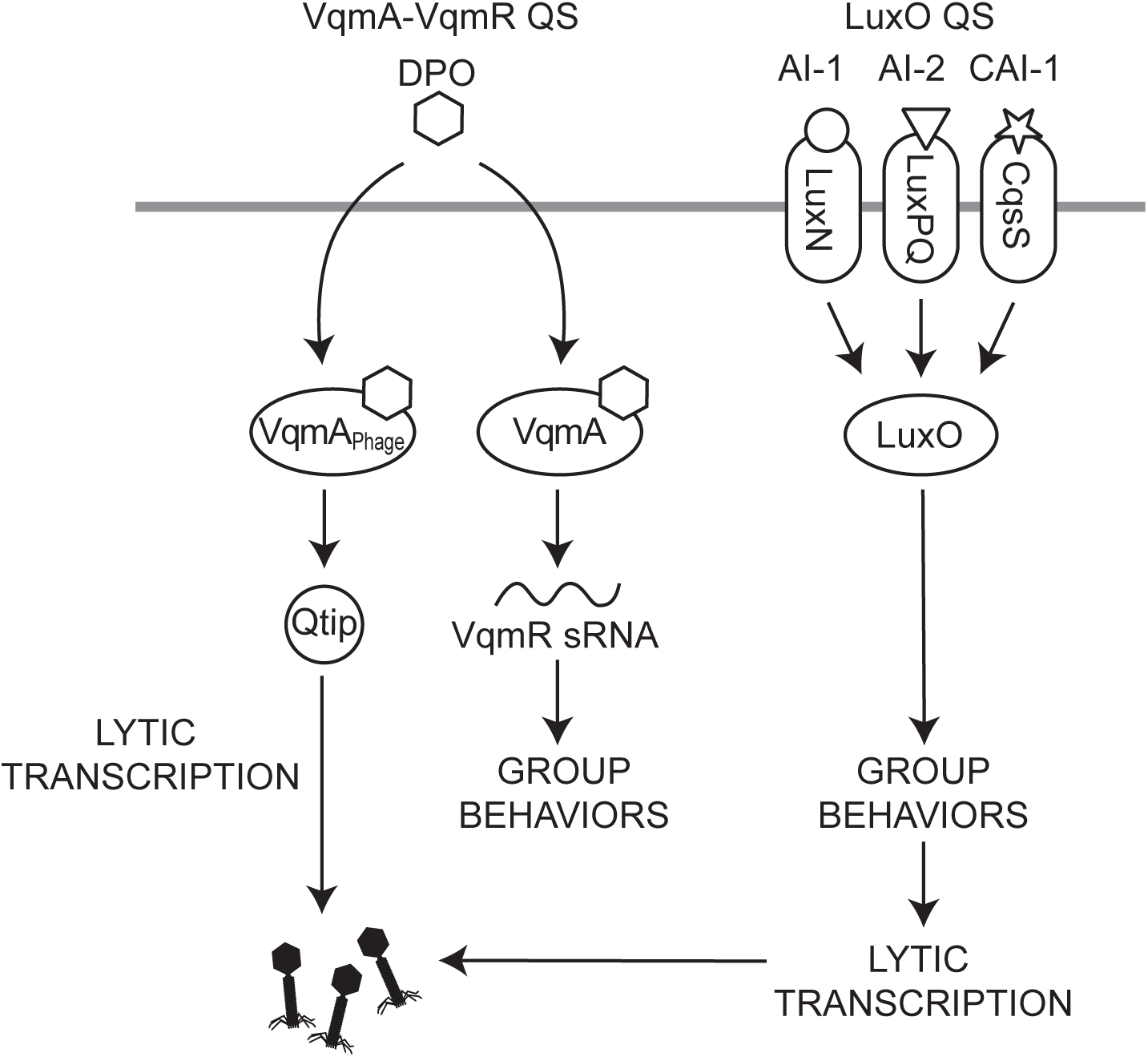
P**h**age **VP882 engages with both host quorum-sensing systems** Both the *V. parahaemolyticus* VqmA-VqmR and LuxO QS systems influence the life cycle of phage VP882 by both promoting lytic replication at high host cell densities. See text for details.

The life cycle of phage VP882 is influenced by both the *V. parahaemolyticus* VqmA-VqmR^19^ and LuxO QS systems^6,20^, both promoting lytic replication at high host cell densities. Specifically, the phage VP882-encoded VqmA_Phage_ receptor binds DPO^21^ and DPO-VqmA_Phage_ activates transcription of *qtip* encoding Qtip, a small protein anti-repressor that sequesters and inactivates cI, the phage VP882 lytic repressor^13^. Via an undefined mechanism, at high cell density, signaling through the LuxO QS system increases phage VP882 lytic gene transcription^20^ favoring lytic replication over lysogeny^23^.

If or how phage VP882 gauges the susceptibility of potential new host cells and whether hommoimmunity or superinfection exclusion occurs have not been investigated. Toward an understanding of those features, here we define factors that influence phage VP882 transmission. We identify the receptor of phage VP882 as the O3:K6 K-antigen that is specific to its host strain. We show that phage access to the O3:K6 K-antigen is controlled by LuxO-directed QS regulation of a transport system responsible for export of polysaccharides that shield the K-antigen. QS-mediated derepression of the genes encoding the transport system at high cell density prevents phage VP882 adsorption and infection. We demonstrate that phage VP882 does not encode a superinfection exclusion mechanism, so *V. parahaemolyticus* cells that are lysogenic for phage VP882 can be infected by virions released from neighboring cells. Following superinfection, DNA from newly infecting phages can recombine with the resident phage genome resulting in superlysogenic conversion. We propose that QS-driven host superinfection via reliance on a specific K-antigen receptor could foster phage VP882 genomic diversification in lysogenic *V. parahaemolyticus* populations.

## Results

### The O3:K6 K-antigen is the phage VP882 receptor in *V. parahaemolyticus*

A primary determinant of phage transmission concerns the ability of phage to attach to host cells. In our continuing effort to define mechanisms that enable phage VP882 to propagate and to connect those mechanisms to its capability to surveil host QS, we investigated phage VP882 host range by identifying the *V. parahaemolyticus* phage receptor that enables infection. To do this, we performed transposon mutagenesis on *V. parahaemolyticus* 882, the natural host of phage VP882, that had been cured of the phage. We infected the pool of transposon insertion mutants with phage VP882 and isolated colonies that were not killed (Figure 2A). The notion was that surviving mutants were resistant to phage VP882 infection. Complicating this simple premise is that phage VP882 is a temperate phage, so infection can result in either lytic replication or lysogenization. Thus, following mutagenesis and infection, some host cells that survived could have been lysogens. To overcome this issue, the strain we used for the transposon mutagenesis carried a plasmid harboring arabinose-inducible *q* (denoted pBAD-*q*)^23^. Q is the phage VP882 activator of lytic gene expression. Consequently, the addition of arabinose to the surviving cells induced the lytic program in lysogens, thereby killing those cells. Presumably, this strategy resulted in survival of only *V. parahaemolyticus* 882 mutants resistant to phage VP882 infection (Figure 2A).

**Figure 2:**
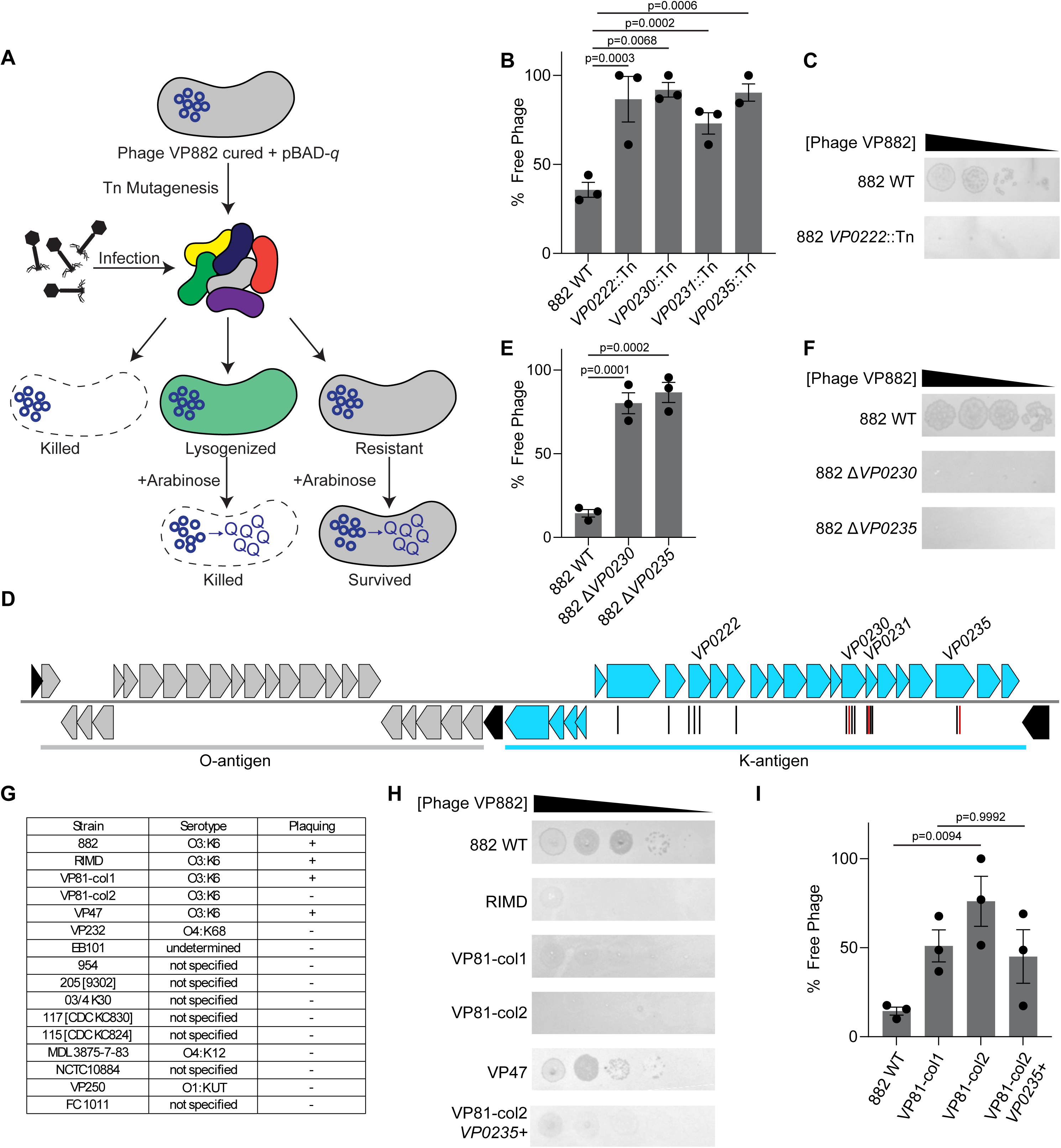
T**h**e **O3:K6 K-antigen is the phage VP882 receptor in *V. parahaemolyticus*** A) Diagram of transposon screen to identify genes encoding the phage VP882 receptor. B) Quantitation of phage VP882 adsorption. Bacteria were incubated with phage VP882 lysate (final MOI 10^-3^-10^-4^) for 20 min. Bacteria and adsorbed phage particles were separated from unabsorbed (free) phage by centrifugation. Free phage were quantified by plaque assay on *V. parahaemolyticus* cured of phage VP882. Percent free phage was calculated relative to a sample lacking bacteria, and the value from that experiment was set to 100%. C) Phage VP882 plaque formation on the designated strains. Ten-fold serial dilutions of phage VP882 lysate (starting at 10^-8^-10^-9^ PFU/mL) were spotted on lawns of the indicated *V. parahaemolyticus* strains embedded in 0.5% LB top agar overlaid onto 1.5% LB agar. Plaques were quantified after 16-20 h of incubation at 37 °C. D) Diagram of *V. parahaemolyticus* 882 O- and K-antigen loci. The K-antigen is encoded by *VP0215*-*VP0237* (blue). Transposon insertion sites are depicted as vertical lines below genes. Black and red indicate insertions on the + and – strands, respectively. Genes shown in black are markers that commonly flank O- and K-antigen loci in *V. parahaemolyticus* (from left to right *coaD*, *gmhD*, and *rjg*). E and F) As in B and C, respectively, for the designated strains. G) Table reporting *V. parahaemolyticus* serotypes, when known. + indicates plaques detected in 3 biological replicates. - indicates no plaques detected. H) As in C for the designated strains. col1 and col2 denote the translucent and opaque morphotypes, respectively. *VP0235+* indicates repair of the frameshift in *VP0235*. I) As in B for the designated strains. In B, E, and I, data are presented as means of 3 biological replicates and error bars denote SEMs. Significance was determined by one-way ANOVA with Tukey’s test for multiple comparisons to establish adjusted p-values. In C, F, and H, a representative image from 3 biological replicates is shown.

We screened surviving colonies for phage VP882 genes using PCR to ensure that the isolated mutants were phage-free. All phage-free mutants had acquired a translucent colony morphology that differed from that of the *V. parahaemolyticus* 882 parent, suggesting a modification to the cell surface (Figure S1A). Indeed, phage VP882 could not adsorb to nor form plaques on these mutants (Figure 2B and C, respectively; panel C shows one representative mutant). We sequenced the transposon insertion sites in 19 of these mutants. All of the insertions mapped to the K-antigen/capsule locus (*VP0215-VP0237*)^24^ suggesting that this surface polymer could be the phage VP882 receptor (Figure 2D, Table S1). We generated in-frame deletions of *VP0230* (encoding a glycosyltransferase) and *VP0235* (encoding an epimerase) in the *V. parahaemolyticus* 882 chromosome and verified by adsorption and plaque assays that the phage could not adsorb nor propagate (Figure 2E and F, respectively), and identical to the original transposon insertion mutants, the deletion strains displayed the translucent colony morphology (Figure S1B; Δ*VP0235* is shown as the example). Some phages can use multiple host receptors to infect. To test whether phage VP882 might have such a capability, we assessed whether lysogenization of the *V. parahaemolyticus* Δ*VP0235* mutant occurred after extended incubation with a variant of phage VP882 carrying chloramphenicol resistance (denoted VP882*^gp^*^38::Cm^). *gp38* encodes a phage VP882 accessory gene^25^ and elimination of it does not affect phage VP882 lysis of host cells (Figure S1C). This approach yielded no chloramphenicol resistant lysogens indicating that the *V. parahaemolyticus* K-antigen is the receptor for phage VP882 (Figure S1D).

Our identification of the K-antigen as the determinant of phage VP882 infectivity suggested that phage VP882 may specifically infect *V. parahaemolyticus* strains of the O3:K6 serotype, the serotype of its *V. parahaemolyticus* 882 host. To test this possibility, we screened our collection of *V. parahaemolyticus* isolates for susceptibility to phage VP882 infection by plaque assay. While not all of the serotypes of our isolates have been reported, phage VP882 formed plaques exclusively on isolates classified as serotype O3:K6 (Figure 2G and H; we discuss the differences in the extent of plaquing in the next section). Verifying this finding, only *V. parahaemolyticus* O3:K6 isolates produced chloramphenicol resistant colonies following infection with phage VP882*^gp^*^38::Cm^ (Figure S1E).

Notably, in the case of one of our O3:K6 serotype isolates, *V. parahaemolyticus* strain VP81, two colony morphologies were present in the stock obtained from ATCC (Figure S1F, designated col1 and col2 for colony type 1 and colony type 2). Colony type 1 had an opaque morphology resembling wildtype (WT) *V. parahaemolyticus* 882, and this *V. parahaemolyticus* VP81 morphotype was susceptible to phage VP882 infection and underwent lysis. By contrast, colony type 2 was translucent, resembling our *V. parahaemolyticus* 882 K-antigen mutants and was resistant to phage VP882 plaquing (Figure 2H and I). Whole genome sequencing of one colony of each morphotype revealed that the *V. parahaemolyticus* VP81 translucent colony 2 type possesses a frameshift mutation in *VP0235*, a gene in the K-antigen locus and a gene also identified in our transposon mutagenesis. Repair of this point mutation converted the colony from translucent to opaque (Figure S1F) and, moreover, restored the ability of phage VP882 to adsorb and plaque (Figure 2I and H, respectively). Together, our data support the model that the O3:K6 K-antigen is the receptor for phage VP882, and phage VP882 specifically attaches to and infects *V. parahaemolyticus* O3:K6 isolates because of its dependence on this antigen.

The particular features of *V. parahaemolyticus* O- and K-antigens that correspond to distinct serotypes have not been defined, so we cannot predict what K6 K-antigen moieties are recognized by phage VP882 nor how its relationship to the O3 O-antigen appendage affects phage VP882 recognition. Nonetheless, to preliminarily probe the basis for specificity of phage VP882 host range, we analyzed the O- and K-antigen loci in the genome sequences of our collection of *V. parahaemolyticus* strains. In agreement with our observations, all isolates designated O3:K6 harbor highly similar O- and K-antigen loci whereas these loci are markedly different in the genomes of non-O3:K6 strains. Few genes encoded in the K-antigen locus of *V. parahaemolyticus* O3:K6 can be identified in the other *V. parahaemolyticus* strains suggesting that their K-antigens are distinct. We conclude that the O3:K6 K-antigen is the essential receptor used by phage VP882.

### LuxO-driven quorum sensing controls phage VP882 plaquing

Phage VP882 plaqued on all isolates of *V. parahaemolyticus* O3:K6 possessing genes encoding functional K-antigens. However, the extent of plaquing differed depending on the strain (Figure 2H). Specifically, when plated on *V. parahaemolyticus* 882 and *V. parahaemolyticus* VP47, phage VP882 readily formed large plaques at all phage concentrations. By contrast, on *V. parahaemolyticus* RIMD and *V. parahaemolyticus* VP81, phage VP882 formed plaques only at the highest concentration tested and the plaques were small. While *V. parahaemolyticus* VP47 and VP81 have not been well studied, one established difference between *V. parahaemolyticus* 882 and *V. parahaemolyticus* RIMD concerns their QS systems^20^. *V. parahaemolyticus* 882 harbors an 897 base pair deletion that eliminates the *vqmR* promoter, the *vqmR* coding sequence, and a portion of the non-coding region upstream of *vqmA*. Additionally, the *V. parahaemolyticus* 882 *luxO* gene (denoted *luxO_882_*) harbors a 36 bp deletion that causes LuxO_882_ to function as a phospho-mimetic. These mutations lock the respective QS systems into their low cell density modes. Given the connections between host QS and phage VP882 lifestyle transitions, we wondered whether these differences in QS capabilities could influence the differences in phage VP882 plaquing on the various host strains. To probe this possibility, we repaired one or both QS systems in *V. parahaemolyticus* 882 and assessed phage VP882 plaquing (Figure 3A and B). Restoration of VqmA-VqmR QS in *V. parahaemolyticus* 882 (denoted *vqmRA^+^*) did not affect phage VP882 plaquing ability nor plaque size, and that was irrespective of the status of *luxO*. By contrast, when LuxO was restored to WT function (denoted *luxO*^+^), phage VP882 made fewer and smaller plaques, similar to its behavior when plated on *V. parahaemolyticus* RIMD that possesses a functional LuxO (Figures 3A and 2H, respectively). Since plaque assays are performed at high cell density, this result indicated that LuxO-directed QS controls the ability of phage VP882 to plaque on *V. parahaemolyticus.* To verify this supposition, we replaced the native *luxO_882_*with either *luxO^D61E^* or *luxO^D61A^* that, respectively, lock *V. parahaemolyticus* into the low cell density and high cell density QS modes. Phage VP882 plaquing on *V. parahaemolyticus* 882 carrying *luxO^D61E^* was comparable to that on the parent *V. parahaemolyticus* 882 strain that carries the naturally low cell density locked *luxO*^882^ variant. *V. parahaemolyticus* 882 carrying *luxO^D61A^* displayed phage VP882 plaquing comparable to that on strains with WT *luxO* (Figure 3A and B).

**Figure 3:**
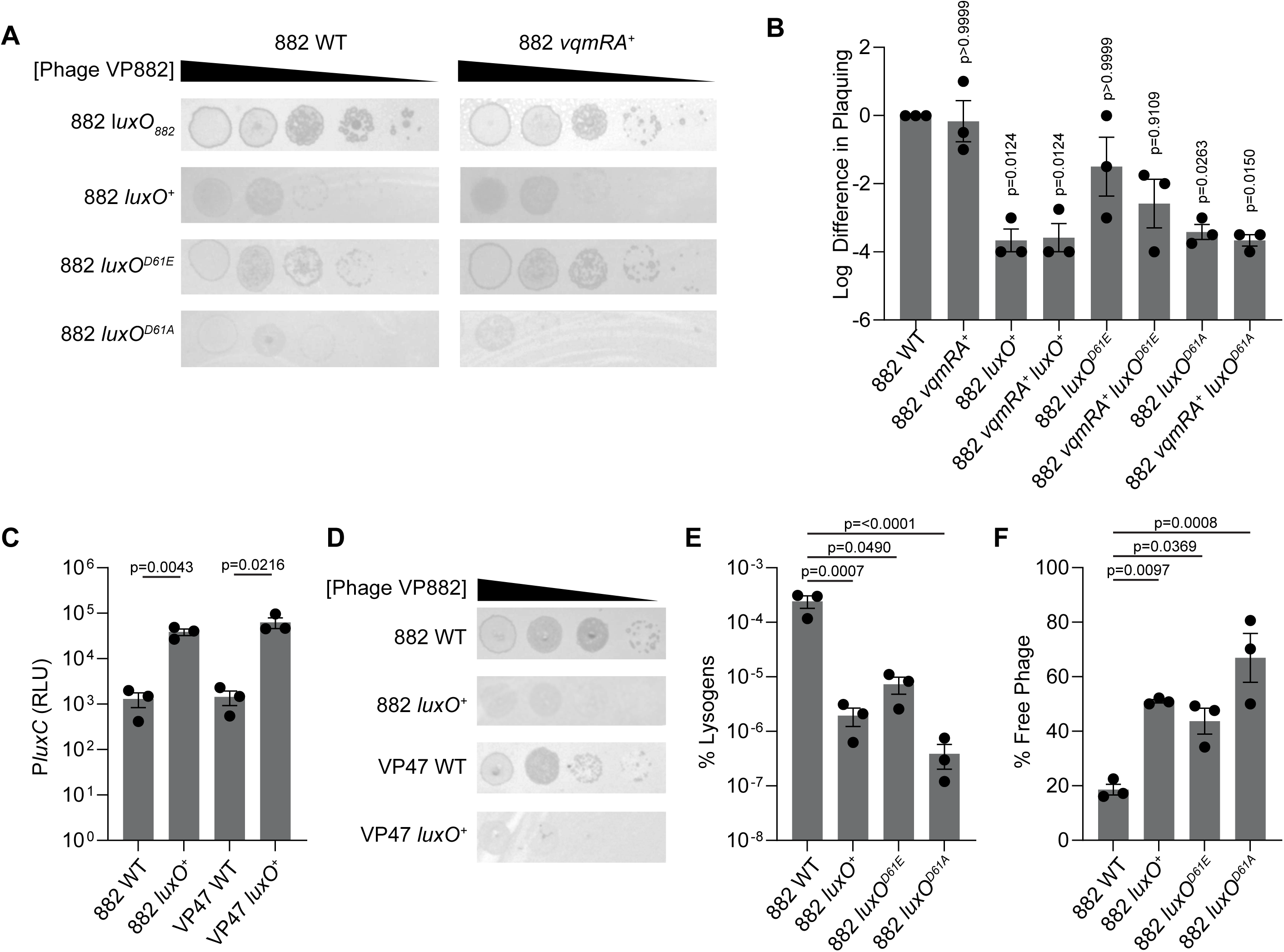
LuxO-driven quorum sensing controls phage VP882 infection A) Phage VP882 plaque formation on the designated phage-free strains, as in Figure 2C. B) Quantification of plaque formation of phage VP882 on the designated strains from panel A. Shown are the log differences in scores for phage VP882 clearance following infection of each strain compared to that for infection on *V. parahaemolyticus* 882 (see Methods). C) Bioluminescence output from P*luxC*-*luxCDABE* after 300 min of growth in LM at 30 °C of the designated strains. Relative light units (RLU) are bioluminescence normalized to OD_600_. D) Phage VP882 plaque formation on the designated strains as in Figure 2C. E) Quantitation of phage VP882*^gp38^*^::Cm^ lysogens following infection of the designated phage-free strains (MOI 10^-5^-10^-6^). F) Quantitation of phage VP882 adsorption as in Figure 2B. In B, C, E, and F, data are presented as means of 3 biological replicates and error bars denote SEMs. Significance was determined by one-way ANOVA with Tukey’s test for multiple comparisons to determine adjusted p-values as indicated. In B, the reported p-values above each bar specify comparisons to *V. parahaemolyticus* 882. In A and D, a representative image from 3 biological replicates is shown.

The plaquing phenotypes associated with the different *luxO* alleles mirrored the genotypes of the tested *V. parahaemolyticus* O3:K6 isolates. As mentioned, LuxO_882_ locks *V. parahaemolyticus* 882 into the low cell density QS state, whereas LuxO in *V. parahaemolyticus* RIMD displays WT function and transitions from the low cell density to the high cell density QS state during growth. LuxO of *V. parahaemolyticus* VP47 harbors an M15R substitution. Analogous to *V. parahaemolyticus* 882 and all *V. parahaemolyticus* strains carrying *luxO^D61E^*, *V. parahaemolyticus* VP47 was incapable of activating a P*luxC* reporter, the canonical QS-activated luciferase promoter that is used as the standard readout for QS function in vibrios^20^ (Figure 3C). Thus, *V. parahaemolyticus* VP47 is locked in low cell density QS mode. Indeed, changing the arginine at position 15 back to methionine in *V. parahaemolyticus* VP47 LuxO restored density-dependent activation of P*luxC* (Figure 3C). Consistent with this finding, phage VP882 plaques only formed on *V. parahaemolyticus* VP47 carrying the repaired LuxO at high phage concentrations, and the plaques were smaller than those on the parent strain with the *luxO^M15R^* allele (Figure 3D). Together, our data show that signaling through the LuxO-controlled QS pathway dictates the robustness of phage VP882 plaquing on *V. parahaemolyticus*.

### *VPA1602-VPA1604* are quorum-sensing-controlled components that shield *V. parahaemolyticus* from infection by phage VP882

We previously reported that LuxO QS signaling inhibits phage VP882 lysogenization at high cell density in *V. parahaemolyticus* RIMD^23^. Here, we show that a similar phenomenon occurs in *V. parahaemolyticus* 882, the natural host for phage VP882: higher numbers of lysogens formed during infection when the host is locked into low cell density QS mode than following infection of the host locked into high cell density QS mode (Figures 3E and S2A). One possibility was that these differences stemmed from differences in the ability of the phage to adsorb to its *V. parahaemolyticus* host when the host exists in the low and high cell density locked QS modes. Indeed, 3-fold less phage VP882 adsorption occurred to *V. parahaemolyticus* 882 encoding WT *luxO* and *luxO^D61A^* than to *V. parahaemolyticus* 882 harboring the *luxO_882_* and *luxO^D61E^*alleles (Figures 3F and S2B). These data indicate that a QS-controlled factor influences phage VP882 entry into naïve host cells. We considered that QS might control production of the K-antigen. However, examination of RNA-sequencing data from *V. parahaemolyticus* 882 carrying either WT *luxO*, *luxO_882_, luxO^D61E^*, or *luxO^D61A^* showed no significant differences in K-antigen transcript levels (Figure S3A)^20^. Thus, we suspected that some other component regulated by LuxO exists in *V. parahaemolyticus* strains and affects phage VP882 adsorption and entry.

To identify the putative factor that inhibits phage VP882 entry into *V. parahaemolyticus* 882, we carried out a selection for *V. parahaemolyticus* 882 mutants displaying enhanced phage VP882 adsorption and entry at high cell density. Specifically, we generated a transposon mutant library in *V. parahaemolyticus* 882 carrying *luxO^D61A^*, infected it with phage VP882*^gp^*^38::Cm^, and selected chloramphenicol resistant lysogens. Sequencing of the transposon insertion sites in 54 lysogens (Table S2) revealed that nearly all insertions resided in unique locations. Most interesting to us were four insertions in or near the *VPA1602-4* operon. *VPA1602-4* are predicted to encode homologs of Wza, Wzb, and Wzc. Wzabc proteins are involved in export of extracellular polysaccharides such as those required for the *Escherichia coli* Group 1 capsules^26^. Deletion of this operon in *V. parahaemolyticus* strains of the O3:K6 serotype have no effect on K-antigen production^24^, so we hypothesized that VPA1602-4 are involved in the production and export of a polysaccharide that blocks access of phage VP882 to the K-antigen. Indeed, deletion of *VPA1602-4* restored the ability of phage VP882 to adsorb to *V. parahaemolyticus* 882 *luxO^D61A^* (Figure 4A). Furthermore, adsorption to all tested *V. parahaemolyticus* 882 strains required the K-antigen including in the absence of *VPA1602-4* (Figure 4A). Consistent with our findings, transcription of *VPA1602-4* is activated at high cell density (Figure S3B and C)^20^ and the homologous operon in *Vibrio harveyi* (*VIBHAR_06665-7*) is repressed at low cell density^27^. These patterns, coupled with our data showing that mutation of *VPA1602-4* promotes phage VP882 adsorption to *V. parahaemolyticus* 882 *luxO^D61A^*, suggest that VPA1602-4 are involved in production and export of a polysaccharide that blocks phage access to the K-antigen.

**Figure 4:**
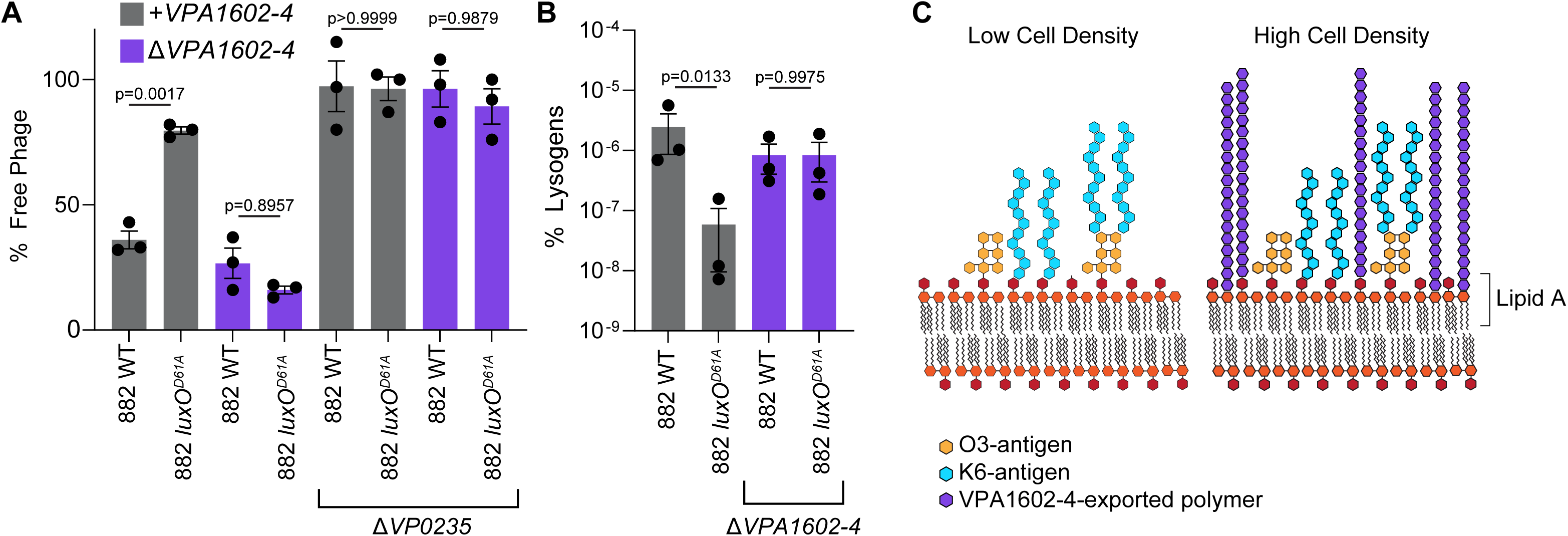
VPA1602-VPA1604 are quorum-sensing-controlled components that shield *V. parahaemolyticus* from infection by phage VP882 A) Quantitation of phage VP882 adsorption as in Figure 2B. B) Quantitation of phage VP882 lysogens as in Figure 3E. Data are presented as means of 3 biological replicates and error bars denote SEMs. Significance was determined by one-way ANOVA with Tukey’s test for multiple comparisons to establish adjusted p-values. C) Diagram of *V. parahaemolyticus* K-antigen and the putative extracellular polysaccharide that shields it.

The export substrates of VPA1602-4 in *V. parahaemolyticus* are undefined. Table S2 shows that, in the above screen, we also isolated singleton transposon insertions in genes involved in polysaccharide biosynthesis pathways (*galU, nagE,* and *neuC*) that enabled phage VP882 entry. We considered that these additional genes could provide clues concerning the polysaccharide produced by VPA1602-4. However, deletion of these genes had no effect on phage VP882 adsorption, so we conclude that they are not required for production of the polysaccharide exported by VPA1602-4 (Figure S3D). We did not investigate the functions of these genes further.

Possibly, the VPA1602-4 system exports multiple substrates that affect K-antigen access to phage VP882 explaining why we did not identify the polymer in the transposon screen.

Deletion of *VPA1602-4* enabled phage VP882 to adsorb to all *V. parahaemolyticus* 882 hosts irrespective of QS genotype (Figures 3A and S3E). Restoration of phage VP882 adsorption via elimination of *VPA1602-4* also rescued the ability of phage VP882 to lysogenize each *V. parahaemolyticus* strain under study to near equivalent levels (Figures 3B and S3F) while having no discernable effect on phage VP882 plaquing (Figure S3G). These results indicate that the major determinant of the LuxO QS effect on phage VP882 lysogenization of *V. parahaemolyticus* 882 is through suppression of phage VP882 virion adsorption to host cells and not via effects on the subsequent lysis-lysogeny decision. Thus, LuxO QS regulation of *VPA1602-4* is a host mechanism that influences the spread of phage VP882 in *V. parahaemolyticus* O3:K6 populations by shielding the K6-antigen (Figure 4C).

### Superinfection of *V. parahaemolyticus* phage VP882 lysogens leads to phage VP882 genome recombination

The above data define how LuxO QS influences phage VP882 transmission to naïve hosts. An outstanding question remained regarding the fate of phage VP882 virions released into a population of QS *V. parahaemolyticus* O3:K6 cells in which virions are likely to encounter hosts that are already lysogens. Superinfection exclusion, via alteration of the expression or function of the phage receptor, is often employed by lysogens to prevent re-infection^12^. We wondered if phage VP882 encodes a superinfection exclusion mechanism that would prevent it from infecting lysogens.

We examined superinfection by plaque assay. Phage VP882 did not form plaques when plated on *V. parahaemolyticus* 882 that was already lysogenic for phage VP882 (Figure 5A). However, in this experiment, homoimmunity, mediated by the resident phage VP882 cI repressor of lysis, could have prevented infecting phages from initiating the lytic replication program necessary for plaque formation even if the virions had adsorbed to and infected lysogens. By exploiting a phage VP882 lysogen that cannot produce functional virions, we found that phage VP882 can indeed adsorb to *V. parahaemolyticus* 882 cells that are already lysogenized with phage VP882 (Figure 5B). Thus, no superinfection exclusion mechanism prevents adsorption of phage VP882 virions to lysogenized hosts.

**Figure 5:**
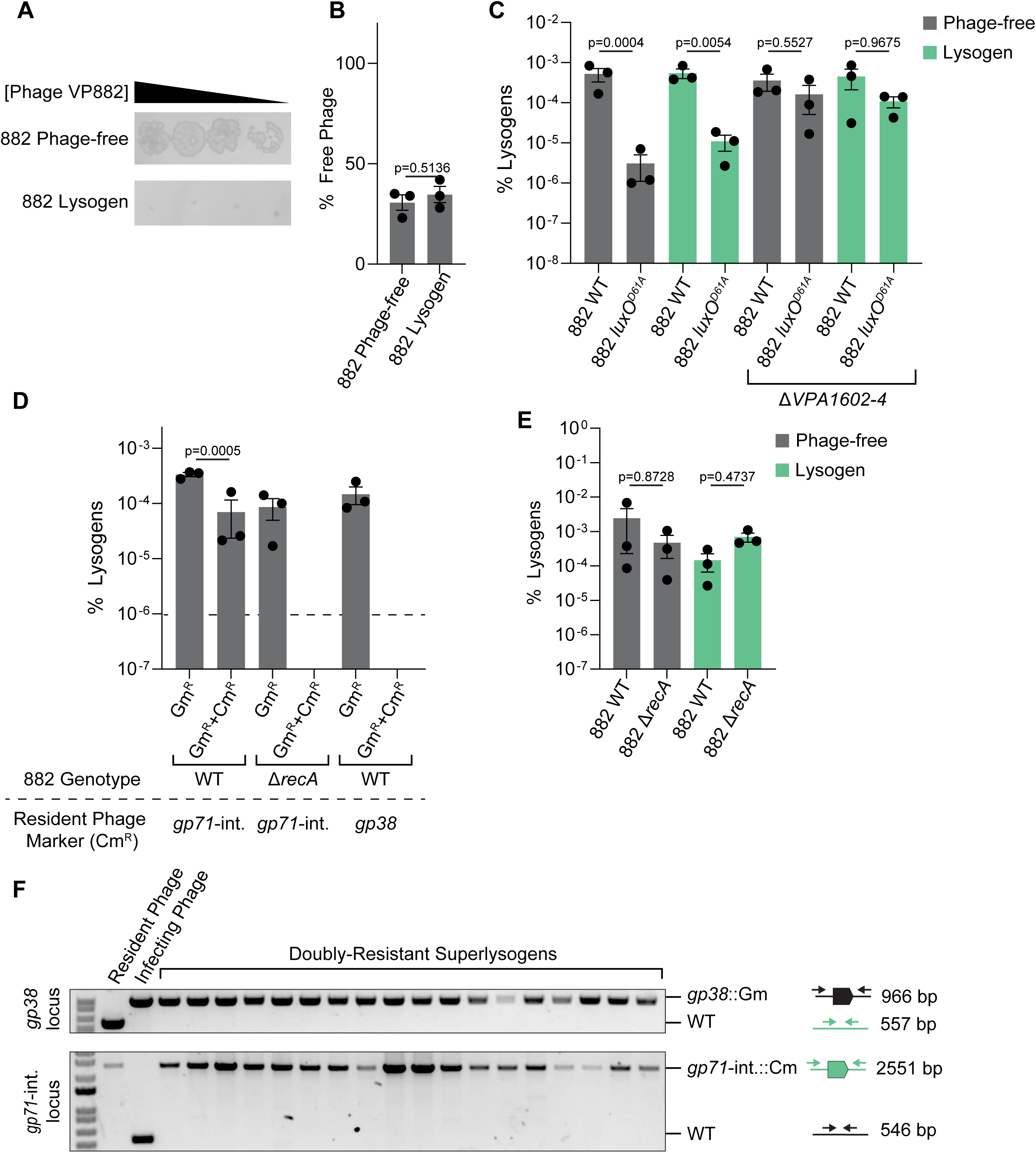
Superinfection of *V. parahaemolyticus* phage VP882 lysogens leads to phage VP882 genome recombination A) Phage VP882 plaque formation as in Figure 2C. A representative image from 3 biological replicates is shown. B) Quantitation of phage VP882 adsorption as in Figure 2B. The phage used for infection, phage VP882^Ctr::Tn*5*^, does not produce functional virions that could interfere with quantitation. C-E) Quantitation of phage VP882 lysogens as in Figure 3E except that in panels D and E, phage VP882*^gp38^*^::Gm^ was the infecting phage. In panel D, lysogens carried phage VP882*^g71-^*^intergenic::Cm^. F) PCR products resulting from amplification of the *gp38* and *gp71*-intergenic loci in the resident and infecting phage VP882 genomes and doubly-resistant superlysogens from the experiment in Figure 5D. Diagrams depict the expected PCR product size for each locus in the resident phage (VP882*^g71-^*^intergenic::Cm^, green) and the infecting phage (VP882*^gp38^*^::Gm^, black). 12-18 colonies were assayed in 3 biological replicates, and a representative gel is shown. In B, C, D, and E, data are presented as means of 3 biological replicates and error bars denote SEMs. In B, significance was determined by an unpaired t-test. In C, D, and E significance was determined by one-way ANOVA with Tukey’s test for multiple comparisons to establish adjusted p-values.

Following adsorption to *V. parahaemolyticus* 882 lysogens, we wondered whether injection of the phage genome occurred. To probe this issue, we assessed whether infection of *V. parahaemolyticus* 882 lysogens with phage VP882*^gp^*^38::Cm^ generated chloramphenicol resistant colonies. Indeed, chloramphenicol resistant lysogens were produced at levels similar to those resulting from infection of *V. parahaemolyticus* 882 that had been cured of phage VP882 (Figure 5C). Thus, phage VP882 virions can both attach to lysogenic cells and inject their genomes. We call new lysogens, which originate from existing *V. parahaemolyticus* 882 phage VP882 lysogens and acquire DNA from an infecting phage VP882, superlysogens.

The influence of QS on phage VP882 lysogenization of *V. parahaemolyticus* 882 via regulation of *VPA1602-4* was maintained during superinfection. Specifically, lysogens of *V. parahaemolyticus* 882, which is naturally locked into the low cell density QS mode, were converted to chloramphenicol resistant superlysogens at a higher frequency than were *V. parahaemolyticus* 882 lysogens locked into high cell density QS mode (Figure 5C). Deletion of *VPA1602-4* restored lysogenization of both the phage-cured and lysogenized high cell density locked QS *V. parahaemolyticus* 882 strains to the levels of the corresponding *V. parahaemolyticus* 882 low cell density locked cured and lysogenic strains (Figure 5C). These results indicate that LuxO regulation of *VPA1602-4* occurs in *V. parahaemolyticus* 882 lysogens and non-lysogens, and thus, host QS status dictates the outcome of phage VP882 superinfection by affecting phage entry.

The transfer of chloramphenicol resistance following phage VP882*^gp^*^38::Cm^ infection of *V. parahaemolyticus* 882 lysogens indicates that lysogens acquire superinfecting phage DNA. We considered the fates of the resident and superinfecting phage genomes during phage VP882 superinfection and superlysogenization. Phage VP882 is a linear plasmid-phage that exists in multiple copies per cell. Superinfection could result in a heterogeneous population of phage genomes. Alternatively, superinfection could result in exchange of the existing phage genomes with the infecting phage genome. Finally, superinfection could result in recombination of resident and infecting phage genomes. To test which possibility occurs, we infected *V. parahaemolyticus* 882 lysogens with phage VP882 harboring a chloramphenicol marker in an intergenic region near the 3’ terminus of the phage genome (denoted VP882*^gp^*^71^*^-^*^intergenic::Cm^). We investigated whether the chloramphenicol resistance marker was maintained in lysogens that acquired gentamicin resistance during superinfection with phage VP882 carrying a gentamicin resistance marker in *gp38* (denoted VP882*^gp^*^38::Gm^). *gp38* is located near the center of the phage genome (Figure S4). We reasoned that if the phage genomes co-exist following superinfection, all superlysogens would be both gentamicin and chloramphenicol resistant. By contrast, if the phage genomes were exchanged, all superlysogens would exhibit only gentamicin resistance. Third and lastly, if recombination of phage genomes occurred, some superlysogens would be exclusively gentamicin resistant and some would display resistance to both gentamicin and chloramphenicol. Importantly, our method only allows us to detect superlysogenization events that result in the acquisition of gentamicin resistance, we cannot detect events in which gentamicin is acquired and then lost following recombination (Figure S4). The third possibility occurred (Figure 5D) suggesting that recombination occurs between the resident and newly infecting phage genomes. Deletion of *recA*, encoding the recombinase required to initiate homologous recombination, did not affect the level of lysogenization of naïve *V. parahaemolyticus* 882 nor of *V. parahaemolyticus* 882 lysogenized with phage VP882 (Figure 5E). However, *V. parahaemolyticus* 882 Δ*recA* lysogens carrying VP882*^gp^*^71^*^-^*^intergenic::Cm^ did not yield doubly antibiotic-resistant superlysogens when infected with phage VP882*^gp^*^38::Gm^ (Figure 5D); all of the colonies that had acquired gentamicin resistance had lost chloramphenicol resistance. Furthermore, infection of *V. parahaemolyticus* 882 lysogens carrying VP882*^gp38^*^::Cm^ with phage VP882*^gp^*^38::Gm^ did not yield doubly antibiotic-resistant lysogens (Figure 5D). Thus, when recombination that places both antibiotic resistance markers onto the same phage genome is prevented or is not possible, the superlysogens acquire the entire genome of the infecting phage. Together, these results suggest that only one variant of the phage VP882 genome is maintained in *V. parahaemolyticus* and, during superinfection, recombination occurs between the infecting and resident phage genomes to enable this outcome.

To confirm the above findings, we tested whether all the phage genomes in a *V. parahaemolyticus* 882 superlysogen were identical or not. To do this, we used PCR amplification of the loci containing the resistance cassettes in doubly-antibiotic resistant *V. parahaemolyticus* 882 superlysogens (Figure S4). The logic is if only one phage genome is present, PCR amplification spanning either the *gp38* or the *gp71*-intergenic locus will yield a single PCR product, while if two phage genomes are maintained, PCR amplification should yield two PCR products corresponding to the allele containing and lacking the resistance marker. PCR amplification yielded one product for each locus, and the product size corresponded to the phage with the resistance cassette in each locus (Figure 5F). Additionally, both gentamicin and chloramphenicol resistance were maintained when these superlysogens were grown in the absence of antibiotic selection. This result indicated that the markers were linked and could not independently segregate over multiple generations. These data support our model that recombination occurs between the infecting and resident phage genomes during superinfection and only one variant of the phage genome is maintained.

## Discussion

Phage VP882 engages with both QS systems of its host *V. parahaemolyticus* and funnels the information it garners into control of transitions between lysogeny and lytic replication (Figure 1). Monitoring host QS enables phage VP882 to favor transitioning from lysogeny to lysis at high host cell densities. Presumably, skewing transitions in this direction increases the frequency of released phage VP882 virions encountering new host cells. An outstanding question was whether those encounters are productive given that monitoring host QS cues does not provide the phage information concerning whether or not nearby host cells are already lysogens. Irrespective of encounter frequency, populations of hosts that are predominantly lysogens could restrict phage VP882 transmission if resident VP882 phages deploy homoimmunity or superinfection exclusion mechanisms that block subsequent phage infection or propagation.

Here, we identify and characterize host components that influence transmission of phage VP882 in QS populations of *V. parahaemolyticus*. We identify the receptor used by phage VP882 as the O3:K6 K-antigen that is specific to its host strain (Figure 2B-F). We show that phage access to the O3:K6 K-antigen is controlled by LuxO-driven QS regulation of a transport system that exports polysaccharides that shield the K-antigen from the phage at high cell density (Figure 4A and C). We demonstrate that phage VP882 does not encode a superinfection exclusion mechanism, so *V. parahaemolyticus* lysogens can be superinfected by virions released from neighboring cells (Figure 5D). Our data demonstrate that following superinfection, DNA from newly infecting phage genomes can be maintained in the entirety or can recombine with the resident phage genome resulting in superlysogenic conversion (Figure 5D and F). Thus, phage VP882 avoids unproductive encounters with already lysogenized hosts. We propose that, together, superinfection and phage genome recombination promote phage VP882 spread and genomic diversity in lysogenic *V. parahaemolyticus* populations.

Phage VP882 could employ the specific K-antigen receptor to ensure transmission to similar hosts to which the phage arrives well-adapted. As a plasmid-phage, and we have experimentally demonstrated this trait previously^6^, phage VP882 can persist in, replicate in, and lyse a wide variety of very distantly related/unrelated bacterial species because its RepA module is sufficient for plasmid maintenance and does not require a specific site in the host genome for the phage to lysogenize. However, the narrow distribution of the O3:K6 receptor guarantees that, in natural settings, phage VP882 exclusively infects *V. parahaemolyticus* strains of that serotype. Interestingly, both the LuxO and VqmR-VqmA systems that influence phage VP882 lifestyle transitions are only present in *vibrio spp*^5,6^. Rigid receptor-specificity could serve as a gate-keeping mechanism to ensure the prophage receives the benefits of detecting and responding to the multiple host QS signals by restricting infection to only similar *V. parahaemolyticus* strains. Recently, VP882-like plasmid-phages have been discovered that reside in other bacterial species^25^. It will be fascinating to learn if they too use specific receptors that constrain host range to hosts to which they are likewise highly adapted.

While QS plays an overarching role in interactions between phage VP882 and its *V. parahaemolyticus* host, here, we find that QS also drives host adaptation to the phage. Specifically, at high cell density, LuxO-QS activates production of a polysaccharide export system that protects *V. parahaemolyticus* from phage VP882 adsorption and therefore, infection. In other phage-bacterial systems, QS-controlled restriction of phage propagation is mediated by alterations in phage receptor availability, via upregulation of CRISPR systems, and through control of abortive infection mechanisms^28–31^. In the current case, we find that changes in phage VP882 adsorption do not alter phage VP882 lytic propagation but, rather, affect the ability of phage VP882 to lysogenize its host. Clearly, additional QS-controlled mechanisms that modulate phage VP882 activity exist in *V. parahaemolyticus* that, for example, contribute to differences in phage VP882 plaquing when phage VP882 adsorption is equivalent among strains (Figure S3G). Potentially, *V. parahaemolyticus* has evolved multiple strategies to counteract phage VP882 in response to the phage’s pervasive infective capability.

Rather than deploying a superinfection exclusion mechanism that diverts phage VP882 progeny to infection of naïve hosts^12^ or a mechanism that detects the abundance of lysogens or like phages in the vicinal community^3^, phage VP882 infects lysogens that already harbor phage VP882. While phage VP882 does not propagate immediately following superinfection, as noted above, these infection events are not unproductive, as the newly infecting phage DNA can be maintained in the superinfected host. We propose that coupling surveillance of QS cues with superinfection is especially beneficial to phage VP882 compared to phages that do not possess one or both of these capabilities. First, lysis primarily occurs when many potential host cells are present and infection or superinfection promotes phage transmission. Second, superlysogenization could promote phage VP882 genetic diversity. Our data indicate that the primary outcome of superlysogenization is that a single phage genome is maintained in the host. Consistent with our notion that this process fosters genomic diversity, we demonstrate that, in many cases, superinfection results in recombination between the resident and infecting phage genomes. We detect these events in 1-5% of superlysogens (Figure 5D), but the frequency of recombination is likely higher as only a subset of recombination events can be detected by our markers. Superinfection resulting in either genome exchange or recombination potentially enhances the spread of phage VP882 variants because genomes of variants with reduced fitness can be replaced by genomes or portions of genomes from successfully replicating and successfully superinfecting phages. The narrow host specificity phage VP882 displays via dependence on the K6 antigen presumably increases the likelihood that, if virions encounter a lysogen that the virion can infect, the resident phage genome will be similar enough to the newly infecting phage genome for recombination to occur.

Together, our data reveal that phage VP882 transmission is influenced by both host and phage features that hinge on bacterial QS inputs. At high host cell density, phage VP882 lytic replication is stimulated by inputs from both QS systems of *V. parahaemolyticus* host cells^6,20^. Superinfection enables phage VP882 to overcome the challenge temperate phages face when they induce their lytic cycle in populations of lysogens. In this case, superinfection moreover ensures that phage VP882 can capitalize on its interception of bacterial QS cues. *V. parahaemolyticus* host cells “fight back” by restricting phage VP882 spread using QS-directed production of a receptor-shielding molecule, also at high cell density. Potentially, this host strategy could broadly defend against phage infection. In the context of our current work, the receptor-shielding mechanism tips phage VP882 lysogeny toward naïve host populations and conditions of low host abundance, both of which are non-optimal for phage VP882 to propagate and diversify. Thus, bacterial QS signaling underlies both host and phage behaviors that shape the outcomes of phage VP882 replication and host encounters. These dynamics could differ for other phage-host pairs depending on the selection pressures germane to their specific environments and the mechanisms connecting phage processes to host QS signaling.

## Materials and Methods

### Bacterial strains, reagents, and growth conditions

#### *Escherichia coli* strains were grown with aeration in Lysogeny Broth (LB-Miller, BD-Difco) at 37

°C. *V*. *parahaemolyticus* strains were grown with aeration in LM (LB with 2% NaCl) at 30 °C. Strains used in the study are listed in Table S3. Unless otherwise noted, antibiotics were used at: 50 μg/mL kanamycin (GoldBio), 50 μg/mL polymyxin B (Sigma), 10 μg/mL chloramphenicol (Sigma), and 30 μg/mL gentamicin (Sigma). Diaminopimelic acid (DAP, Sigma) was used at 100 μM. For phage lysis and bioluminescent reporter assays, overnight cultures of *V*. *parahaemolyticus* strains were diluted 1:100 into fresh LM containing antibiotics when appropriate. After 2 h of growth at 30 °C with aeration, cultures were diluted 1:100 into fresh medium with appropriate antibiotics prior to being dispensed (200 μL/well) into 96 well plates (Corning Costar 3903). When needed, 0.05 μg/mL mitomycin C was added at the second dilution step. A BioTek Synergy Neo2 Multi-Mode reader was used to measure OD_600_ and bioluminescence. Relative Light Units (RLU) are bioluminescence divided by OD_600_. P*luxC*-*luxCDABE* expression was measured at 300 min of growth. P*VPA1602*-*luxCDABE* expression was measured at 500 min of growth.

### Cloning techniques

All primers used for plasmid construction and qPCR, listed in Table S4, were obtained from Integrated DNA Technologies. FastCloning^32^or Gibson Assembly (NEB) were employed for plasmid assembly. Briefly, PCR was performed with Q5 polymerase (NEB) to generate cloning inserts. In the case of pTOX2^33^, linear plasmid backbone was generated by restriction digestion with SwaI (NEB). Inserts in assembled plasmids were verified by sequencing (Azenta Genewiz or Plasmidsuarus). Plasmids used in this study are listed in Table S5. Transfer of plasmids into *V*. *parahaemolyticus* was accomplished by conjugation with *E. coli* S17 or MFD*pir*^34^ (a DAP auxotroph) followed by selective plating on LB plates containing appropriate antibiotics. Mutants were generated by allelic exchange with 15% sucrose (pRE112 backbone) or 2% rhamnose (pTOX2 backbone) for counter selection. Mutations were verified by colony PCR or Sanger sequencing. To insert antibiotic cassettes into the phage VP882 genome, PCR products containing the antibiotic cassette flanked by homologous phage VP882 DNA sequences were introduced by natural transformation. Competence of *V. parahaemolyticus* 882 was induced by expression of *tfoX*^35^. Phages carrying antibiotic cassettes were isolated by lysogenization of *V. parahaemolyticus* 882 that had previously been cured of phage VP882.

### Generation of phage lysates

Overnight cultures of *V. parahaemolyticus* 882 strains lysogenized with phage VP882 were diluted 1:100 in LM and incubated for 2 h at 30 °C with aeration (180 rpm). Mitomycin C was added and growth continued for 2.5 h. Lysates of lytic phage VP882 were generated by infection of *V. parahaemolyticus* 882 that had been cured of phage VP882 for 16 h at 30 °C with aeration. Debris was removed from lysed cultures by centrifugation (10,000 x g for 1 min). Clarified supernatant was treated with 0.5 % chloroform for at least 15 min prior to use. Phage lysate titers were quantified by plaque assay.

### Phage VP882 plaque assay

Phage VP882 plaquing on *V. parahaemolyticus* strains was quantified by the double-layer overlay plaque method^36^. *V. parahaemolyticus* was grown from a single colony to stationary phase (16 h). The culture was diluted 1:100 into molten 0.5% LB agar containing 10 mM CaCl_2_. The mixture was poured over 1.5% LB agar in a petri plate and allowed to solidify at room temperature for 10-15 min. 0.1% Evan’s Blue (Sigma) and 0.2% uranine (Sigma) were incorporated into the bottom agar to enhance plaque visibility. 10-fold serial dilutions of phage VP882 lysate (initial titer 10^-8^-10^-9^ PFU/mL) were spotted onto the plate (3.5 µL/spot) and allowed to dry prior to incubation overnight at 37 °C. Plaques were imaged with an IQ800 imager (Cytvia) using the UV setting and black tray with 0.1 s exposure. Because phage VP882 does not form individual plaques on *V. parahaemolyticus* in high cell density QS mode, plaquing was assessed based on clearance and the sizes of plaques as described previously^37^. The lowest 10-fold dilution spot in which clearance occurred was recorded. The log differences in plaquing shown in Figures 3A and S2A are the differences between the plaquing score for the parent *V. parahaemolyticus* 882 strain cured of phage VP882 and the designated strain.

### Phage VP882 adsorption assay

*V. parahaemolyticus* was grown from a single colony to stationary phase (16 h). The culture was adjusted to OD_600_ 2.5 with LB. 50 µL of this culture was combined with 150 µL of a suspension containing 10 mM CaCl_2_ and lysate from a spontaneously occurring lytic phage VP882 variant (final MOI 10^-3^-10^-4^). The mixture was incubated at room temperature for 20 min. Cells and adsorbed phage particles were separated from free phage by centrifugation (10,000 x g for 1 min). 30 µL of the free phage solution was added to 4 mL of 0.5% LB agar containing 10 mM CaCl_2_ and a 1:100 dilution of a stationary phase culture of *V. parahaemolyticus* 882 that had been cured of phage VP882. The mixture was poured into 3 cm petri dishes (Falcon). Plaques were quantified after overnight incubation at 37 °C. The fraction of unadsorbed phage (% free phage) was calculated by comparing the number of phage to that in an identically treated sample lacking bacteria.

### Phage VP882 lysogenization assay

*V. parahaemolyticus* was grown from a single colony to stationary phase (16 h). The culture was adjusted to OD_600_ 2.5 with LB. The cell suspension was mixed at a 1:4 ratio with a suspension containing LB, 10 mM CaCl_2_, and lysate from phage VP882 marked with an antibiotic cassette at an MOI 10^-5^-10^-6^. The mixture was incubated without shaking at 30 °C for 1 h prior to plating on nonselective medium to quantify total cells and on selective medium to quantify lysogens. For experiments to test lysogenization of *V. parahaemolyticus* lacking the O3:K6 K-antigen receptor, the entire lysogenization mixture was diluted into 3 mL of fresh LM containing 10 mM CaCl_2_ and grown for 16 h at 30 °C with aeration before plating on selective medium. Plates were incubated at 37 °C for 16-20 h and colonies counted. Percent lysogens is the number of colonies on selective medium divided by the number on nonselective medium. In all lysogenization quantitation experiments except those examining phage recombination, phage VP882*^gp38^*^::Cm^ was used. In experiments examining phage recombination, lysogens harboring phage VP882*^gp71-^*^intergenic::Cm^ were infected with phage VP882*^gp38^*^::Gm^ lysate. Lysogens resistant to both gentamicin and chloramphenicol were isolated and purified to verify dual-resistance. Homogeneity of each locus was assessed by colony PCR with DreamTaq Polymerase (Thermo). PCR products were analyzed on 1% agarose 1X TAE gels stained with SYBR Safe (Edvotek) and imaged with an IQ800 imager (Cytvia) using the Cy2 setting and black tray with auto-exposure.

### Construction of transposon mutant libraries

Transposon insertion libraries in *V. parahaemolyticus* 882 or *V. parahaemolyticus* 882 *luxO^D61A^* both of which had been cured of phage VP882 were generated with the Mariner transposon on pSC189 using kanamycin selection^38^. *E. coli* MFD*pir*/pSC189 was used as the donor to deliver pSC189 by conjugation. 300 µL of donor and 100 µL of recipient were mixed. The suspension was subjected to centrifugation (8,000 x g for 2 min) and 390 µL of the clarified culture fluid was removed. Cells were resuspended in the residual medium and spotted on 0.45 μm mixed cellulose ester membrane filters (Millipore) that had been placed onto LB agar plates followed by incubation at 37 °C for 2 h. The filters were transferred to microcentrifuge tubes containing 1 mL of PBS and subjected to vortex to dislodge cells. Various dilutions of the cell suspensions were plated on nonselective and selective media to quantify the number of exconjugants. 10 separate matings were performed to generate the library. Each mating yielded ∼35,000-70,000 exconjugants. Nested arbitrary PCR (Neo-F + Arb 1, pSC189-1^st^-3^rd^ + Arb2)^39^ was used to assess randomness of transposition. Exconjugants were collected from the plates by scraping all colonies into LB broth containing appropriate antibiotics, resulting in cell suspensions of OD_600_ of ∼50. Aliquots of the library were frozen in 30% glycerol.

### Transposon mutagenesis screen to identify genes encoding the phage VP882 receptor

pBAD-*q* was introduced into the phage-cured *V. parahaemolyticus* 882 transposon mutants from the above library by conjugation. Exconjugants were selected on LB agar containing chloramphenicol and kanamycin. Selection of survivors of phage VP882 infection was performed by diluting the library of mutants containing pBAD-*q* 1:2,000 or 1:20,000 into 8 separate 10 mL aliquots of LB containing 10 mM CaCl_2_ and phage VP882 to yield 8 different ratios of phage: bacteria at MOI 0.001-2 followed by incubation for 16 h. The infected cultures were back-diluted 1:150 into LB broth and grown for 2.5 h at 30 °C with aeration. Subsequently, 0.2% arabinose was added to induce expression of *q* and the cultures grown for 2 h at 30 °C with aeration before plating. Colonies were screened for phage VP882 by PCR for *vqmA_Phage_* (primers oMRS1 + oMRS2). All opaque colonies were positive for phage VP882 indicating that they were lysogens (see main text). Phage-free colonies were isolated, and the transposon insertion sites were identified by arbitrary PCR and Sanger sequencing with the pSC189-1^st^-3^rd^ primer.

### Transposon mutagenesis screen to identify genes encoding factors that prevent phage VP882 infection of *V. parahaemolyticus* 882 at high cell density

A thawed aliquot of the *V. parahaemolyticus* 882 *luxO^D61A^* transposon mutant library was diluted 1:1,000 in 20 mL of LB broth and incubated for 30 min at 30 °C with aeration for resuscitation. The suspension was diluted into 10 tubes containing 10 mM CaCl_2_ and phage VP882*^gp^*^38::Cm^ at MOI 10^-5^-10^-6^, incubated for 3 h with aeration and then plated on LB + chloramphenicol + kanamycin to select lysogens. Plates were incubated for 20 h at 37 °C and then at room temperature for an additional 24 h. Chloramphenicol-resistant colonies were isolated and transposon insertion sites were identified by arbitrary PCR and Sanger sequencing with the pSC189-1^st^-3^rd^ primer.

### Quantitation and statistical analyses

Software used to collect and analyze data generated in this study consisted of: GraphPad Prism 10 for visualization of data, calculation of statistics, and analysis of growth and reporter-based experiments; Gen5 for collection of growth and reporter-based data; Geneious v2026.0 for primer design and sequencing verification; Amersham ImageQuant for imaging of plaque assays and agarose gels. Data are presented as the means ± SEMs. Numbers of technical and independent biological replicates are indicated in the figure legends.

## Acknowledgements

We are grateful to members of the Bassler laboratory for insightful discussions.

## Author Contributions

Outlined the study: M.R.S and B.L.B. Performed experiments: M.R.S Analyzed data: M.R.S and B.L.B. Interpreted data: M.R.S and B.L.B. Wrote manuscript: M.R.S and B.L.B.

## Data and code availability

No sequencing or custom code is included in this manuscript. All plasmids and strains in this study are available from the corresponding author upon request. All data is available upon request.

---

**Figure S1:**
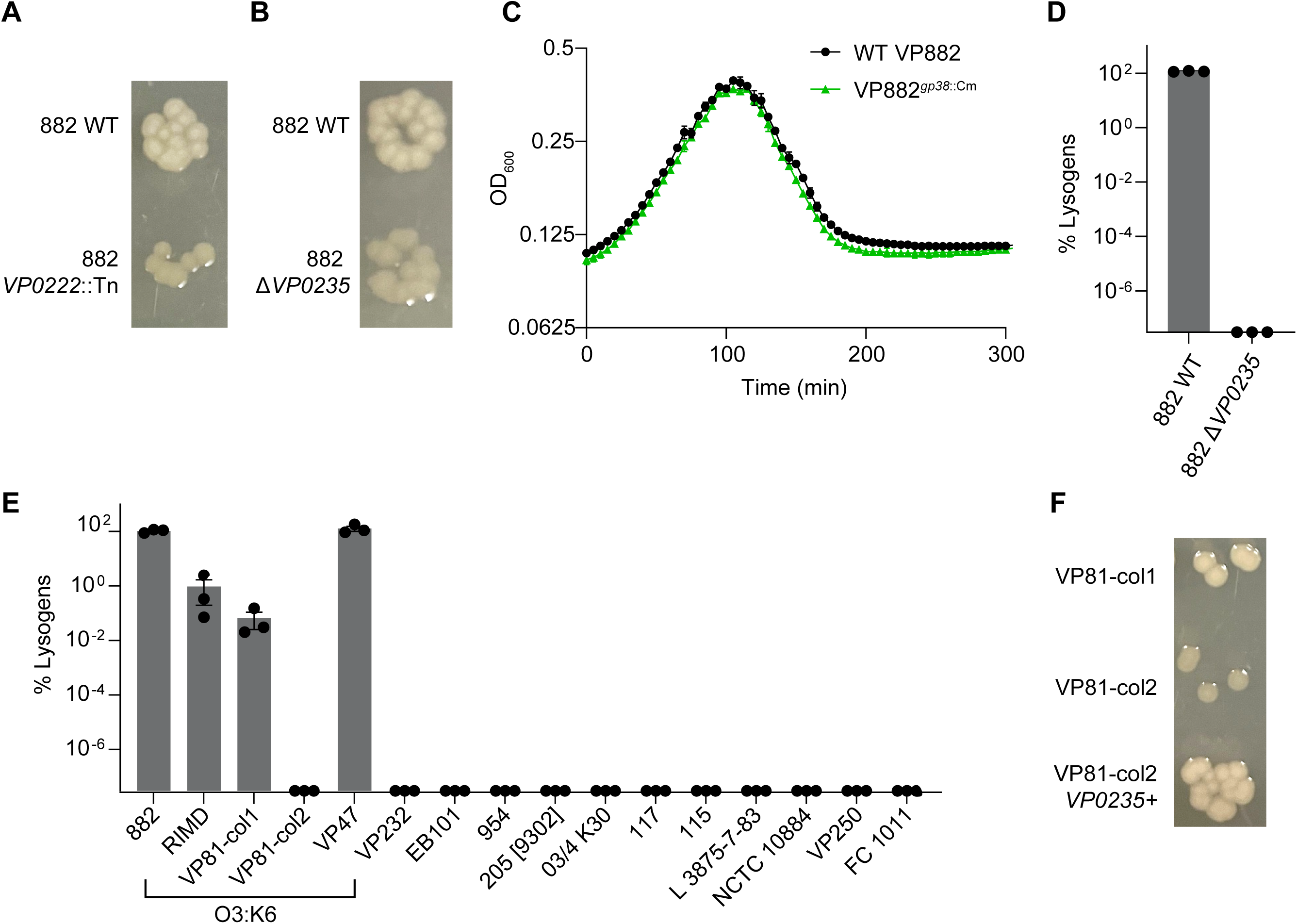
Phage VP882 only infects *V. parahaemolyticus* with the O3:K6 K-antigen. A and B) Colony morphologies of the designated phage-free *V. parahaemolyticus* 882 strains grown on LB agar at 37 °C for 16 h. C) Growth curves of *V. parahaemolyticus* 882 carrying phage VP882 or phage VP882*^gp38^*^::Cm^ treated with mitomycin C (0.05 µg/mL). A representative of 2 biological replicates is shown. Each assay included 2 technical replicates. D) Quantitation of phage VP882*^gp38^*^::Cm^ lysogens following infection of the designated phage-free strains, as in Figure 3E except that lysogenization was carried out for 16 h. E) Quantitation of phage VP882*^gp38^*^::Cm^ lysogens as in Figure 3E. F) Colony morphologies of the designated phage-free strains grown on LB agar at 37 °C for 16 h. In A, B, and F, a representative image from 3 biological replicates is shown. In C, D, and E, data are presented as means of 3 biological replicates and error bars indicate SEMs. We note that in panel C, error bars are obscured by the symbols.

**Figure S2:**
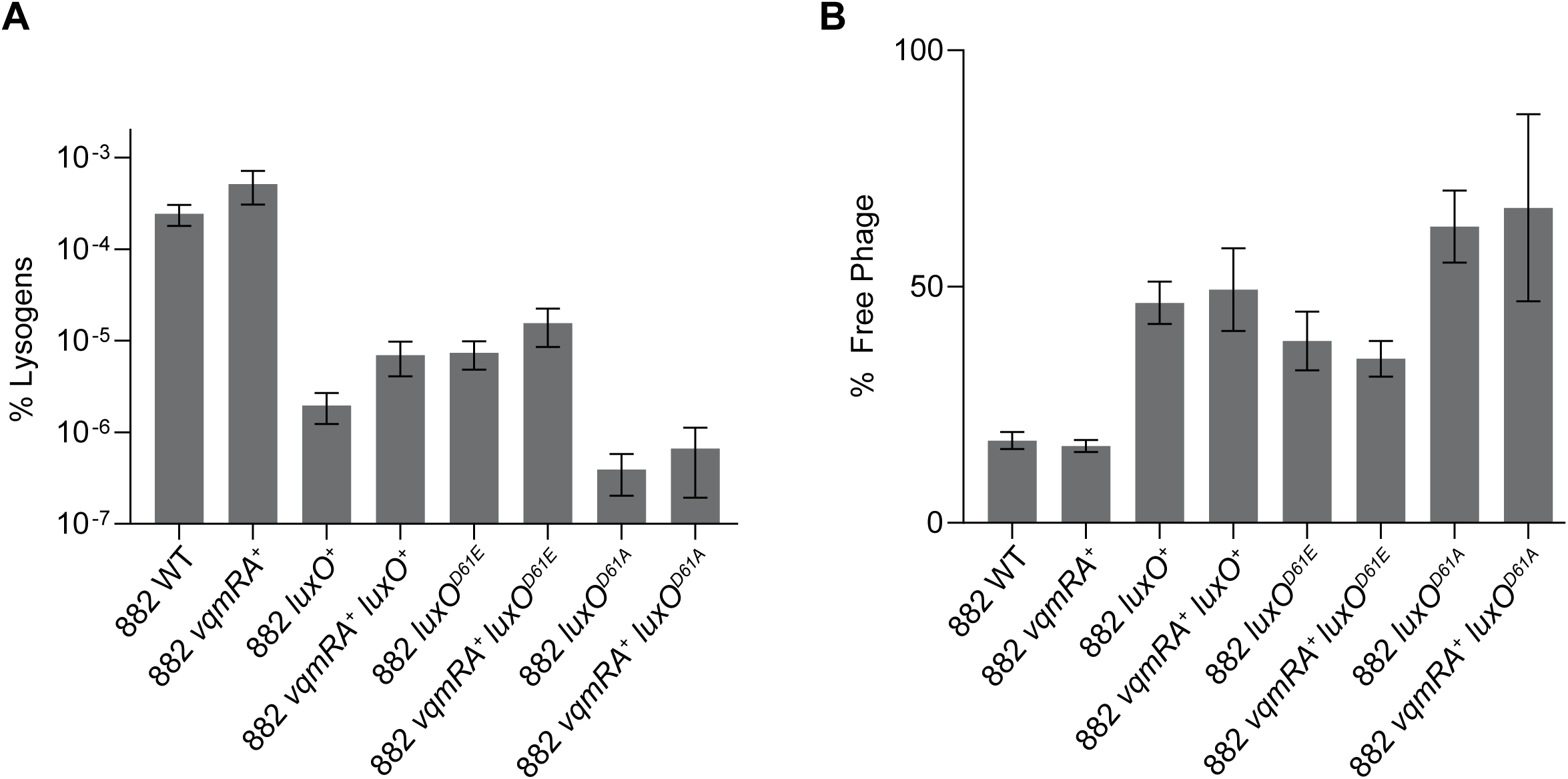
VqmA-VqmR quorum sensing does not affect phage VP882 infection A) Quantitation of phage VP882 lysogens as in Figure 3E. B) Quantitation of phage VP882 adsorption as in Figure 2B. Data are presented as means of 3 biological replicates and error bars indicate SEMs.

**Figure S3:**
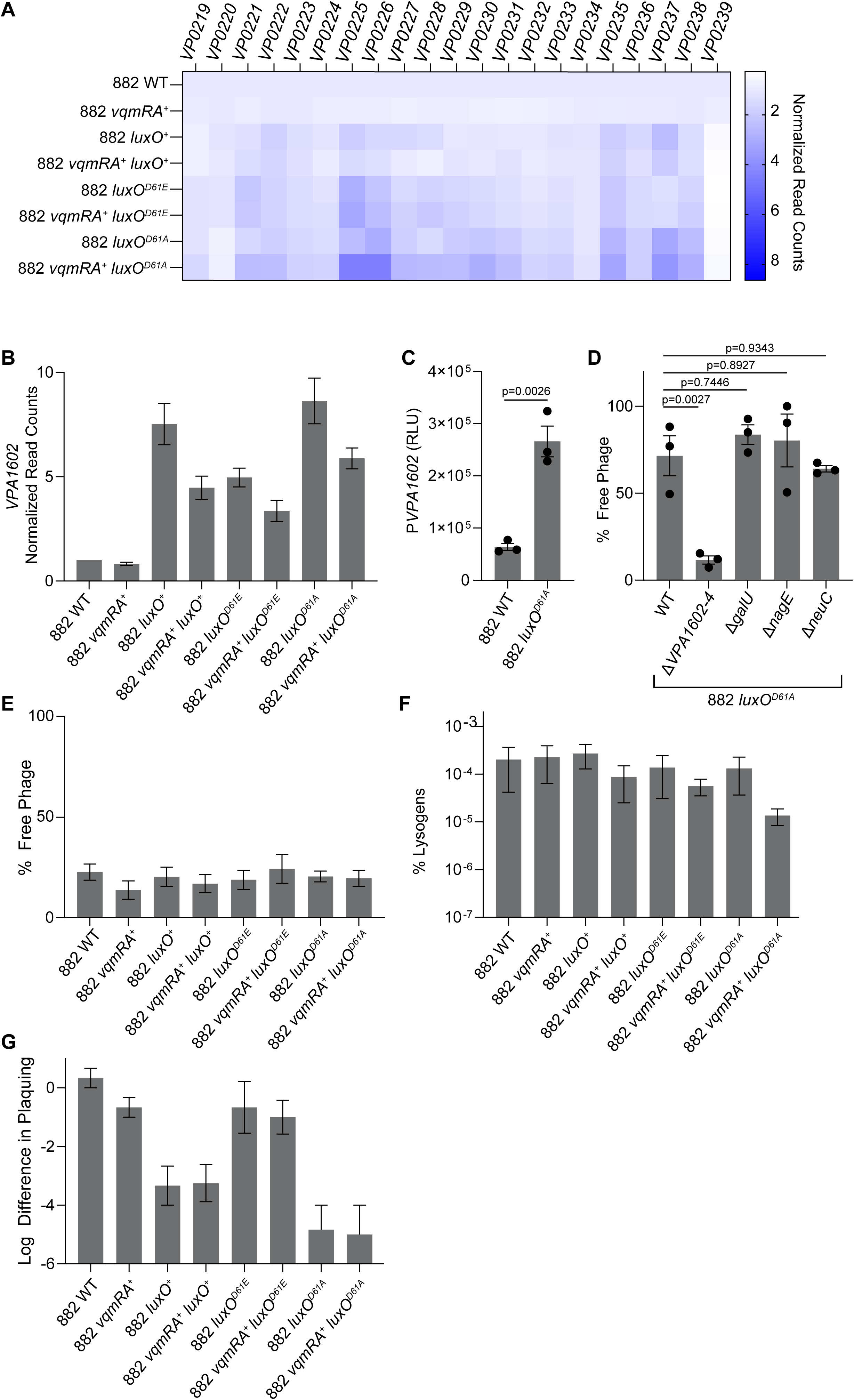
***VPA1602-VPA1604* are controlled by the LuxO quorum-sensing system** A) Heatmap showing quantitation of RNA-seq reads mapping to the *V. parahaemolyticus* K-antigen locus for strains with the indicated QS genotypes after 4 h of growth in LM at 30 °C^20^. The fraction of total reads aligning to each gene were normalized to the corresponding value for *V. parahaemolyticus* 882. B) Quantitation of RNA-seq reads mapping to *VPA1602* from the conditions described in A^20^. C) P*VPA1602*-*luxCDABE* output as in Figure 3C except measurements were taken after 500 min. D and E) Quantitation of phage VP882 adsorption as in Figure 2B. F) Quantitation of phage VP882 lysogens as in Figure 3E. G) Quantification of phage VP882 plaque formation as in Figure 3B. For D-G, data are presented as means of 3 biological replicates and error bars indicate SEMs. In C and D, significance was determined by one-way ANOVA with Tukey’s test for multiple comparisons to establish adjusted p-values.

**Figure S4:**
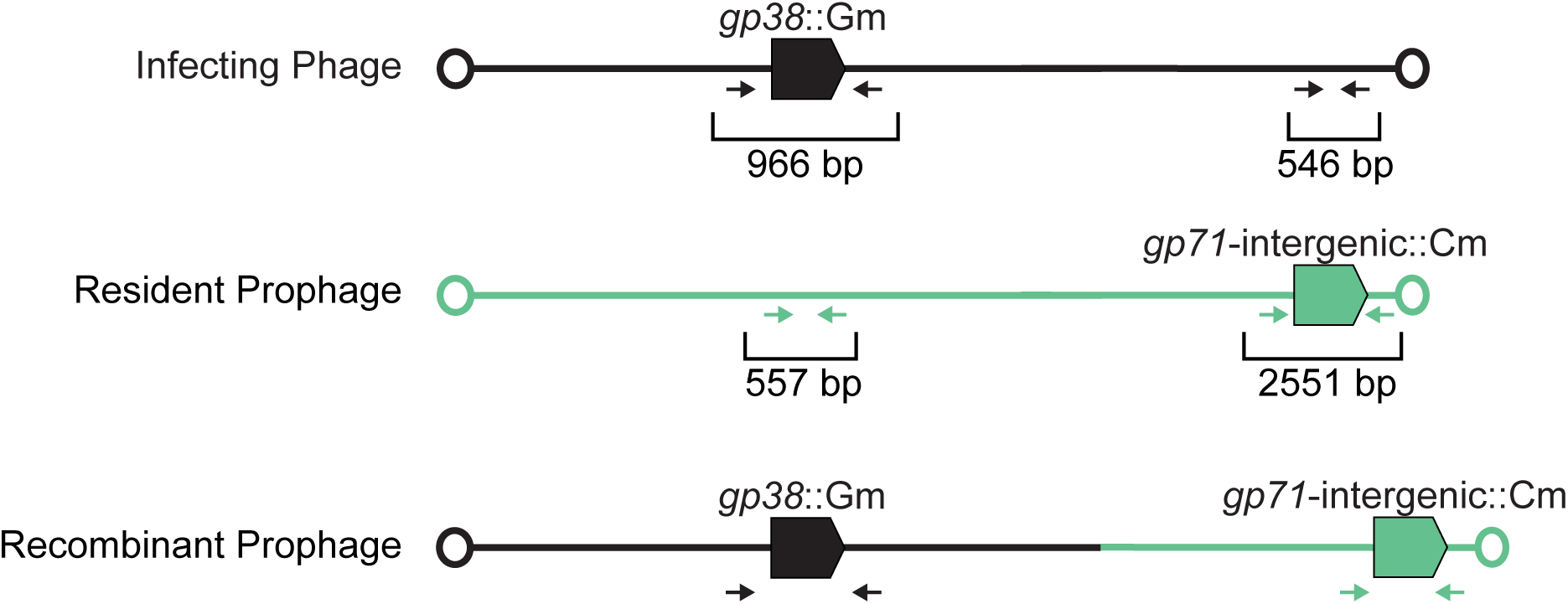
Phage VP882 genome recombination occurs during superlysogenization Diagram of phage VP882 genomes used to monitor recombination in the superinfection experiments reported in Figure 4D and F.

**Table S1:** Transposon insertion sites identified in the screen for the phage VP882 receptor.

| Sequence Number | Insertion Site | Direction | Gene ID |
| --- | --- | --- | --- |
| 1 | 231439 | forward | *VP0220* |
| 2 | 234489 | forward | *VP0221* |
| 3 | 234489 | forward | *VP0221* |
| 4 | 235609 | forward | *VP0222* |
| 5 | 235779 | forward | *VP0222* |
| 6 | 235855 | forward | *VP0222* |
| 7 | 235855 | forward | *VP0222* |
| 8 | 237039 | forward | *VP0224* |
| 9 | 243061 | forward | *VP0230* |
| 10 | 243498 | reverse | *VP0231* |
| 11 | 243643 | forward | *VP0230* |
| 12 | 243914 | forward | *VP0230* |
| 13 | 244073 | forward | *VP0230* |
| 14 | 244219 | forward | *VP0231* |
| 15 | 244285 | forward | *VP0231* |
| 16 | 244285 | forward | *VP0231* |
| 17 | 244479 | forward | *VP0231* |
| 18 | 249145 | forward | *VP0235* |
| 19 | 249360 | reverse | *VP0235* |

**Table S2:** Transposon insertion sites identified in the screen for the factor that prevents phage VP882 adsorption to *V. parahaemolyticus.* Insertions in *VPA1602-4* are shaded gray * indicates the read quality was too poor to identify the insertion site.

| Sequence Number | Insertion Site | Direction | Disrupted locus | Annotation |
| --- | --- | --- | --- | --- |
| Chromosome 1 |  |  |  |  |
| 1 | 77696 | reverse | *VP0068* | *gorA* |
| 2 | 92662 | reverse | *VP0081* | BON Domain protein |
| 3 | 183123 | forward | *VP0168-VP0170*,<br>intergenic | TonB-dependent receptor<br>plug domain-containing<br>protein and ABC<br>transporter |
| 4 | 102615 | reverse | *VP0109* | *engB* |
| 5 | 128809 | reverse | *VP0120* | *glnL* |
| 6 | 196839 | reverse | *VP0183* | Methyl accepting<br>chemotaxis protein |
| 7 | * | forward | *VP0199* | *neuC* |
| 8 | 209620 | forward | 5' *VP0199* | 5' of *neuC* |
| 9 | 210461 | forward | *VP0199* | *neuC* |
| 10 | 253049 | reverse | *VP0238* | MBL fold metallo-<br>hydrolase |
| 11 | 379167 | forward, | 5' *VP0377* | HTH domain protein |
| 12 | 390779 | forward | 3' *VP0388* | methylase |
| 13 | 426994 | forward | *VP0424* | *hldE* |
| 14 | 508389 | forward | *VP0494* | *thrA* |
| 15 | 545647 | reverse | *VP0527* | *nhaR* |
| 16 | 630725 | forward | *VP0605* | rRNA large subunit<br>methyltransferase N |
| 17 | 681151 | reverse | *VP06050*,<br>*VP06051*<br>intergenic | *nadK*, *grpE* |
| 18 | * | forward | *VP0682* | *ribH* |
| 19 | 716483 | forward | *VP0684* | *thiL* |
| 20 | 827688 | forward | 5' *VP0793* | *crr* |
| 21 | * | reverse | *VP0831* | *nagE* |
| 22 | 1022658 | reverse | *VP0980* | DUF2062 domain protein |
| 23 | * | forward | *VP1231* | *fabV* |
| 24 | 1386862 | reverse | *VP1305/VP1306* | *cobU*,<br>adenosylcobinamide-GDP<br>ribazoletransferase |
| 25 | 1421362 | reverse | *VP1340* | Collagenase |
| 26 | 2292011 | reverse | 3' *VP2183* | Response regulator |
| 27 | 2549724 | forward | VP2429-VP2430,<br>intergenic | *radA* |
| 28 | 2643101 | forward | VP2504 | *pcnB* |
| 29 | 2688526 | forward | 5' VP2546 | *csrA* |
| 30 | 2688526 | forward | 5' VP2546 | *csrA* |
| 31 | 2723599 | forward | VP2577 | *rseA* |
| 32 | 2723599 | forward | VP2577 | *rseA* |
| 33 | 2755274 | reverse | VP2611 | *gshB* |
| 34 | 2833881 | forward | VP2683 | PhoH family protein |
| 35 | 2867011 | forward | VP2711 | *galU* |
| 36 | 2971496 | reverse | VP2807 | *rnr* |
| 37 | 3055472 | forward | VP2884 | *dusB* |
| 38 | 3056590 | forward | VP2885 | *fis* |
| 39 | 3056590 | forward | VP2885 | *fis* |
| 40 | 3070488 | reverse | VPr018 | 23S rRNA |
| 41 | 3100523 | forward | VP2911 | 5' of *hupA* |
| 42 | 3220203 | reverse | VP3016 | HvfC/BufC N-terminal<br>domain-containing<br>protein |
| 43 | 3247914 | forward | VP3041 | LysM domain protein |
| Chromosome 2 |  |  |  |  |
| 44 | 37139 | forward | VPA0045 | Ada regulatory protein |
| 45 | 44076 | reverse | VPA0063 | Hypothetical protein |
| 46 | 452925 | reverse | VPA0495 | AraC family transcriptional<br>regulator |
| 47 | * | forward | VPA0627 | 5' of *cyoA* |
| 48 | * | forward | VPA1442 | DUF11 domain protein |
| 48 | 1560611 | forward | VPA1464 | Lcl domain protein |
| 49 | 1700868 | forward | VPA1602 | Polysaccharide export<br>protein Wza |
| 50 | 1700867 | forward | VPA1602 | Polysaccharide export<br>protein Wza |
| 51 | 1701292 | reverse | VPA1602 | Polysaccharide export<br>protein Wza |
| 52 | 1700403 | forward | 5' VPA1602 | 5' of VPA1602<br>Polysaccharide export<br>protein Wza |
| 53 | 1799976 | forward | VPA1678 | *araC* |
| 54 | 2932536 | reverse | VPA2764 | Homoserine<br>dehydrogenase |

**Table S3:**
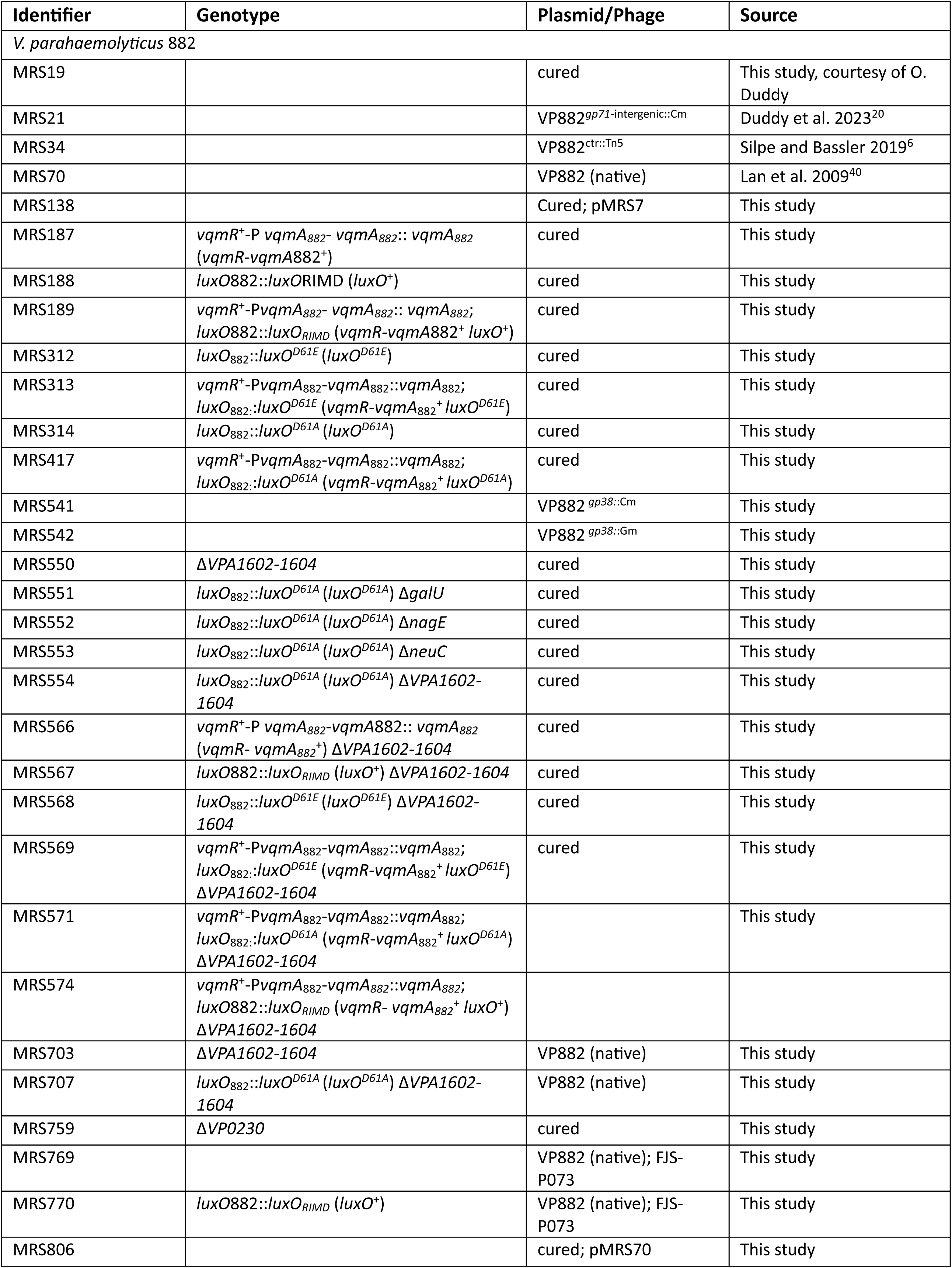
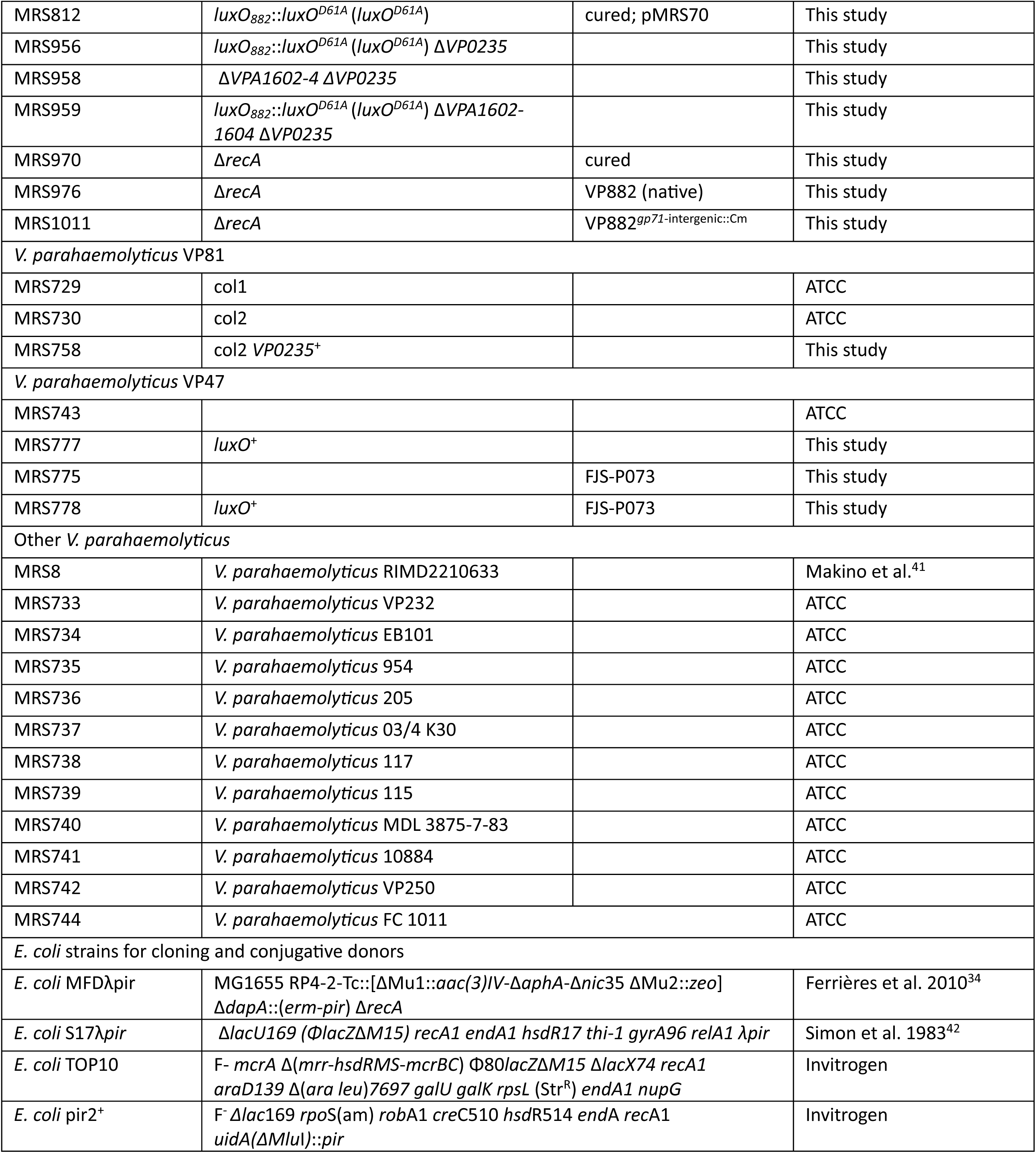
Strains used in this study.

**Table S4:**
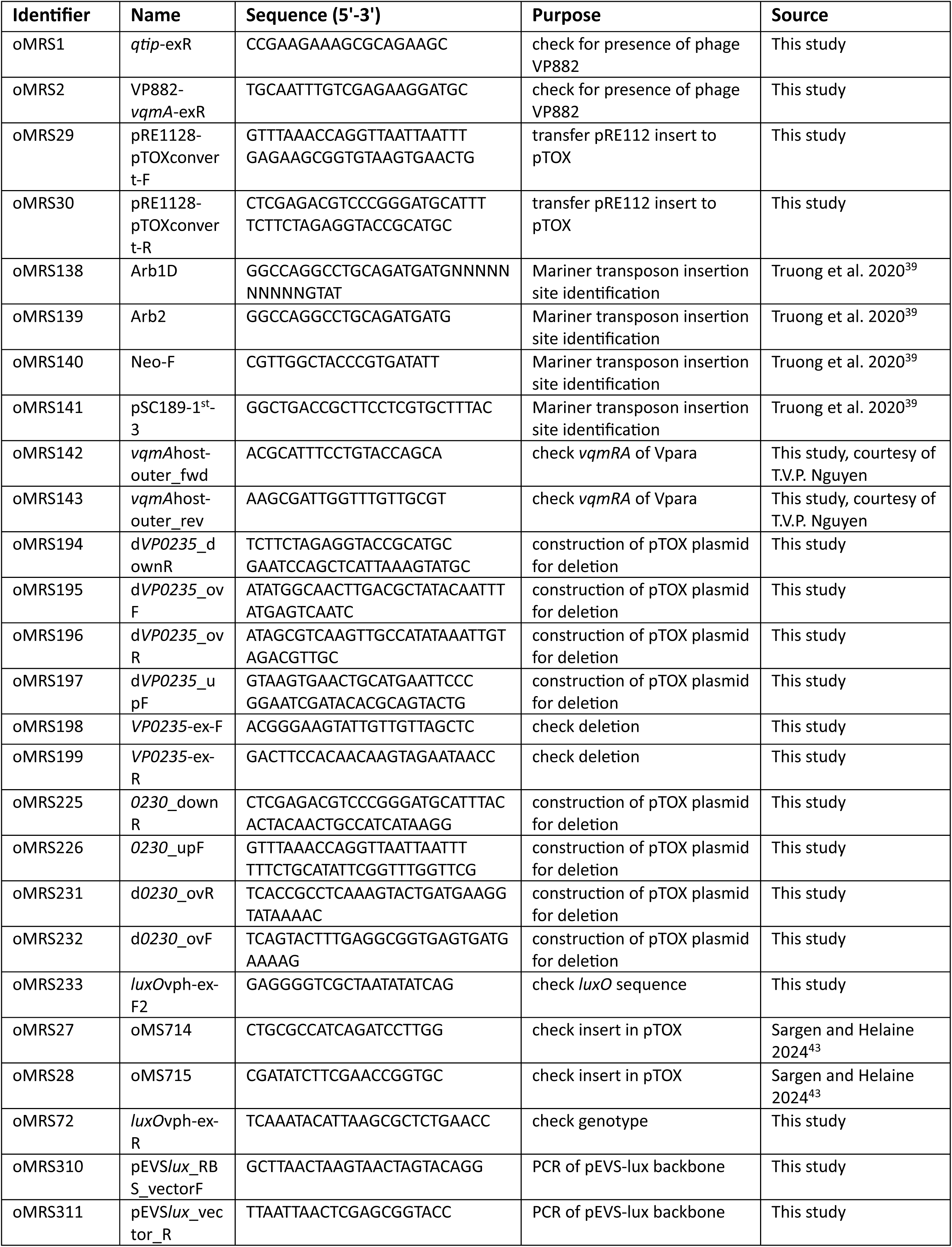
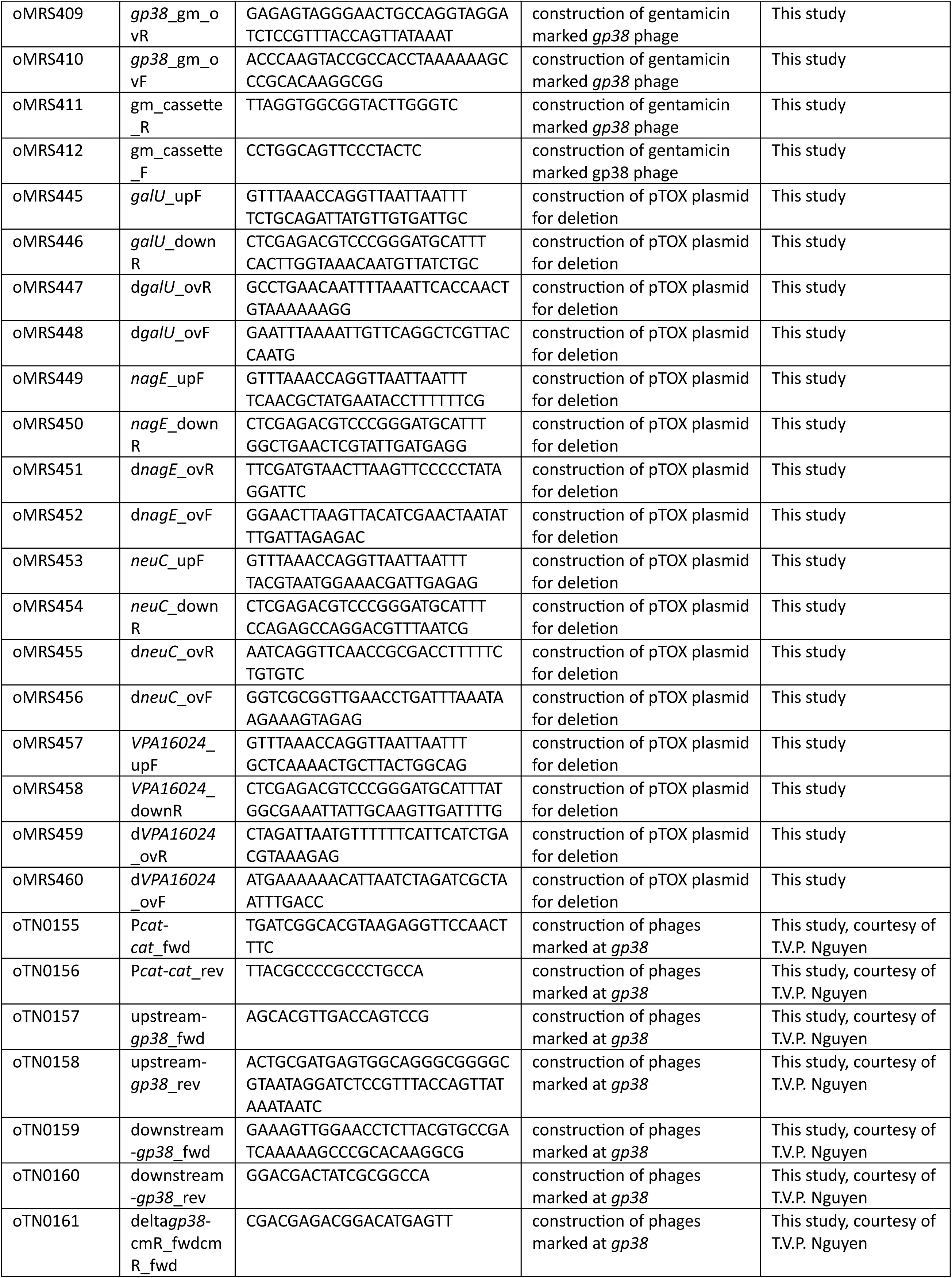
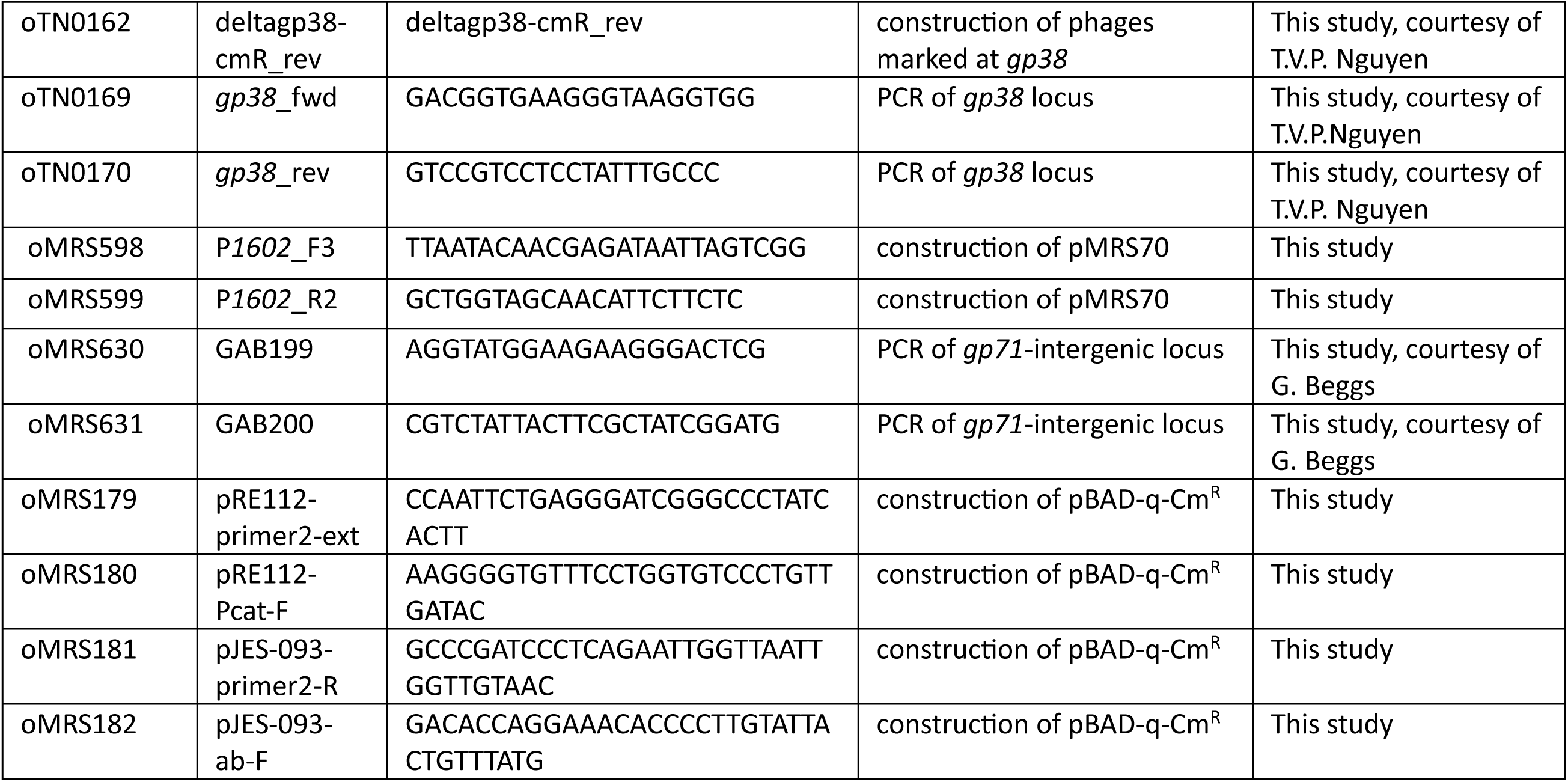
Oligonucleotides used in this study.

**Table S5:**
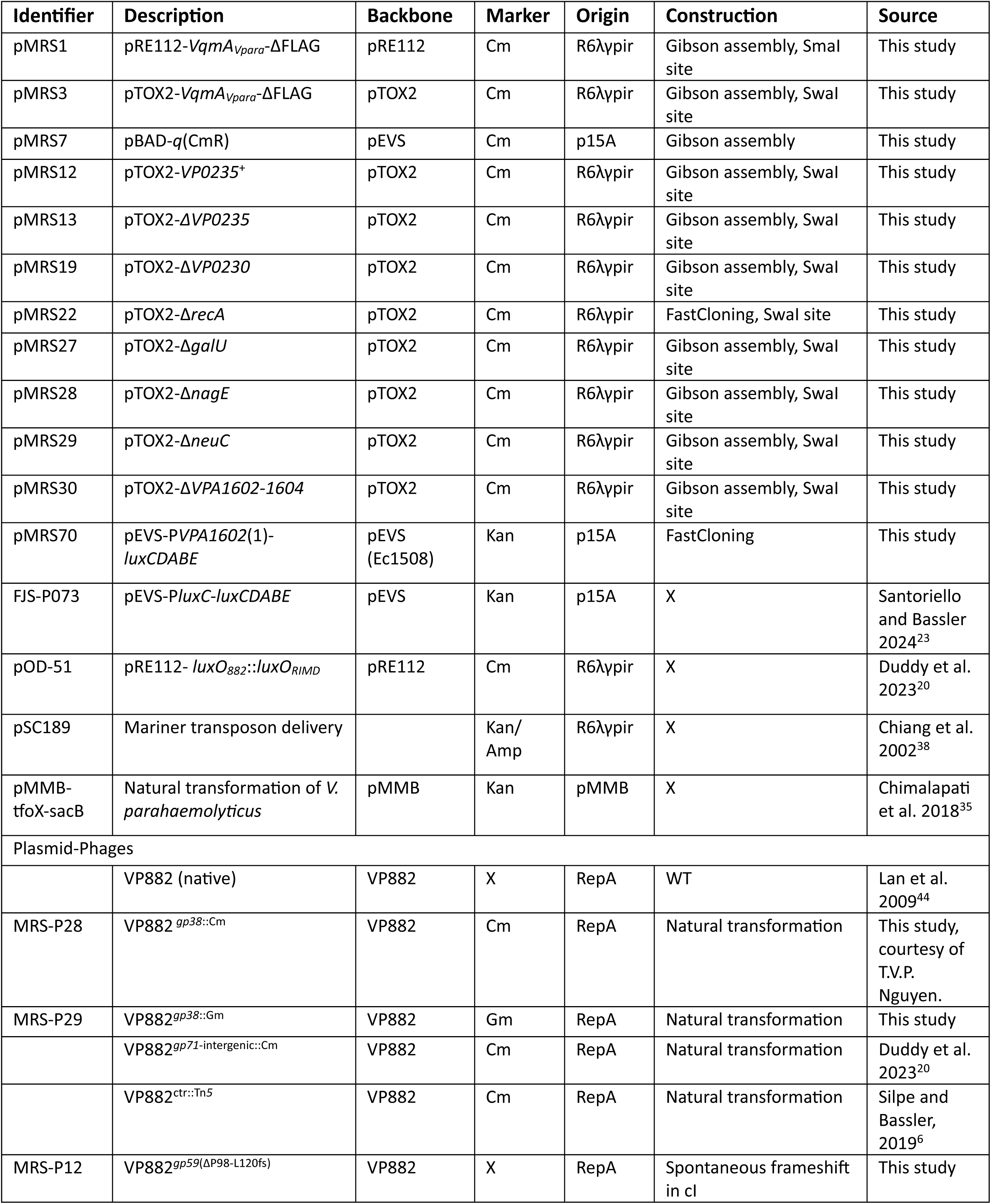
Plasmids used in this study.

## References

1. Howard-Varona, C., Hargreaves, K.R., Abedon, S.T., and Sullivan, M.B. (2017). Lysogeny in nature: mechanisms, impact and ecology of temperate phages. ISME J. 11, 1511–1520. 10.1038/ismej.2017.16.

2. Cheong, K.H., Wen, T., Benler, S., Koh, J.M., and Koonin, E.V. (2022). Alternating lysis and lysogeny is a winning strategy in bacteriophages due to Parrondo’s paradox. Proc. Natl. Acad. Sci. 119, e2115145119. 10.1073/pnas.2115145119.

3. Erez, Z., Steinberger-Levy, I., Shamir, M., Doron, S., Stokar-Avihail, A., Peleg, Y., Melamed, S., Leavitt, A., Savidor, A., Albeck, S., et al. (2017). Communication between viruses guides lysis–lysogeny decisions. Nature 541, 488–493. 10.1038/nature21049.

4. Li, G., Cortez, M.H., Dushoff, J., and Weitz, J.S. (2020). When to be temperate: on the fitness benefits of lysis vs. lysogeny. Virus Evol. 6, veaa042. 10.1093/ve/veaa042.

5. Papenfort, K., and Bassler, B.L. (2016). Quorum sensing signal-response systems in Gram-negative bacteria. Nat. Rev. Microbiol. 14, 576–588. 10.1038/nrmicro.2016.89.

6. Silpe, J.E., and Bassler, B.L. (2019). A host-produced quorum-sensing autoinducer controls a phage lysis-lysogeny decision. Cell 176, 268–280.e13. 10.1016/j.cell.2018.10.059.

7. Silpe, J.E., Duddy, O.P., and Bassler, B.L. (2022). Natural and synthetic inhibitors of a phage-encoded quorum-sensing receptor affect phage–host dynamics in mixed bacterial communities. Proc. Natl. Acad. Sci. 119, e2217813119. 10.1073/pnas.2217813119.

8. Kaiser, A.D., and Jacob, F. (1957). Recombination between related temperate bacteriophages and the genetic control of immunity and prophage localization. Virology 4, 509–521. 10.1016/0042-6822(57)90083-1.

9. Brown, S., Mitarai, N., and Sneppen, K. (2022). Protection of bacteriophage-sensitive Escherichia coli by lysogens. Proc. Natl. Acad. Sci. U. S. A. 119, e2106005119. 10.1073/pnas.2106005119.

10. Łoś, J., Zielińska, S., Krajewska, A., Michalina, Z., Małachowska, A., Kwaśnicka, K., and Łoś, M. (2021). Temperate Phages, Prophages, and Lysogeny. In Bacteriophages, D. R. Harper, S. T. Abedon, B. H. Burrowes, and M. L. McConville, eds. (Springer International Publishing), pp. 119–150. 10.1007/978-3-319-41986-2_3.

11. Bondy-Denomy, J., Qian, J., Westra, E.R., Buckling, A., Guttman, D.S., Davidson, A.R., and Maxwell, K.L. (2016). Prophages mediate defense against phage infection through diverse mechanisms. ISME J. 10, 2854–2866. 10.1038/ismej.2016.79.

12. Bucher, M.J., and Czyż, D.M. (2024). Phage against the machine: the SIE-ence of superinfection exclusion. Viruses 16, 1348. 10.3390/v16091348.

13. Wright, A. (1971). Mechanism of conversion of the *Salmonella* O antigen by bacteriophage ε^34^. J. Bacteriol. 105, 927–936. 10.1128/jb.105.3.927-936.1971.

14. Uc-Mass, A., Loeza, E.J., De La Garza, M., Guarneros, G., Hernández-Sánchez, J., and Kameyama, L. (2004). An orthologue of the cor gene is involved in the exclusion of temperate lambdoid phages. Evidence that Cor inactivates FhuA receptor functions. Virology 329, 425–433. 10.1016/j.virol.2004.09.005.

15. Broadbent, S.E., Davies, M.R., and Van Der Woude, M.W. (2010). Phase variation controls expression of *Salmonella* lipopolysaccharide modification genes by a DNA methylation-dependent mechanism. Mol. Microbiol. 77, 337–353. 10.1111/j.1365-2958.2010.07203.x.

16. Leavitt, J.C., Woodbury, B.M., Gilcrease, E.B., Bridges, C.M., Teschke, C.M., and Casjens, S.R. (2024). Bacteriophage P22 SieA-mediated superinfection exclusion. mBio 15, e02169–23. 10.1128/mbio.02169-23.

17. Taylor, V.L., Patel, P.H., Shah, M., Yusuf, A., Burk, C.M., Sztanko, K.M., Gitai, Z., Davidson, A.R., Koch, M.D., and Maxwell, K.L. (2025). Prophages block cell surface receptors to preserve their viral progeny. Nature 644, 1049–1057. 10.1038/s41586-025-09260-z.

18. Papenfort, K., Silpe, J.E., Schramma, K.R., Cong, J.-P., Seyedsayamdost, M.R., and Bassler, B.L. (2017). A *Vibrio cholerae* autoinducer-receptor pair that controls biofilm formation. Nat. Chem. Biol. 13, 551–557. 10.1038/nchembio.2336.

19. Papenfort, K., Förstner, K.U., Cong, J.-P., Sharma, C.M., and Bassler, B.L. (2015). Differential RNA-seq of *Vibrio cholerae* identifies the VqmR small RNA as a regulator of biofilm formation. Proc. Natl. Acad. Sci. 112. 10.1073/pnas.1500203112.

20. Duddy, O.P., Silpe, J.E., Fei, C., and Bassler, B.L. (2023). Natural silencing of quorum-sensing activity protects *Vibrio parahaemolyticus* from lysis by an autoinducer-detecting phage. PLoS Genet. 19, e1010809. 10.1371/journal.pgen.1010809.

21. Huang, X., Duddy, O.P., Silpe, J.E., Paczkowski, J.E., Cong, J., Henke, B.R., and Bassler, B.L. (2020). Mechanism underlying autoinducer recognition in the *Vibrio cholerae* DPO-VqmA quorum-sensing pathway. J. Biol. Chem. 295, 2916–2931. 10.1074/jbc.RA119.012104.

22. Silpe, J.E., Bridges, A.A., Huang, X., Coronado, D.R., Duddy, O.P., and Bassler, B.L. (2020). Separating functions of the phage-encoded quorum-sensing-activated antirepressor Qtip. Cell Host Microbe 27, 629–641.e4. 10.1016/j.chom.2020.01.024.

23. Santoriello, F.J., and Bassler, B.L. (2024). The LuxO-OpaR quorum-sensing cascade differentially controls Vibriophage VP882 lysis-lysogeny decision making in liquid and on surfaces. PLOS Genet. 20, e1011243. 10.1371/journal.pgen.1011243.

24. Chen, Y., Dai, J., Morris, J.G., and Johnson, J.A. (2010). Genetic analysis of the capsule polysaccharide (K antigen) and exopolysaccharide genes in pandemic Vibrio parahaemolyticus O3:K6. BMC Microbiol. 10, 274. 10.1186/1471-2180-10-274.

25. Santoriello, F.J., and Bassler, B.L. (2026). A family of linear plasmid phages that detect a quorum-sensing autoinducer exists in multiple bacterial species. mBio 17, e02320–25. 10.1128/mbio.02320-25.

26. Whitfield, C. (2009). Structure and assembly of *Escherichia coli* capsules. EcoSal Plus 3, 10.1128/ecosalplus.4.7.3.

27. Shao, Y., Feng, L., Rutherford, S.T., Papenfort, K., and Bassler, B.L. (2013). Functional determinants of the quorum-sensing non-coding RNAs and their roles in target regulation. EMBO J. 32, 2158–2171. 10.1038/emboj.2013.155.

28. Høyland-Kroghsbo, N.M., Mærkedahl, R.B., and Svenningsen, S.L. (2013). A quorum-sensing-induced bacteriophage defense mechanism. mBio 4, e00362–12. 10.1128/mBio.00362-12.

29. Tan, D., Svenningsen, S.L., and Middelboe, M. (2015). Quorum sensing determines the choice of antiphage defense strategy in vibrio anguillarum. mBio 6, e00627. 10.1128/mBio.00627-15.

30. Patterson, A.G., Jackson, S.A., Taylor, C., Evans, G.B., Salmond, G.P.C., Przybilski, R., Staals, R.H.J., and Fineran, P.C. (2016). Quorum sensing controls adaptive immunity through the regulation of multiple CRISPR-Cas systems. Mol. Cell 64, 1102–1108. 10.1016/j.molcel.2016.11.012.

31. Høyland-Kroghsbo, N.M., Paczkowski, J., Mukherjee, S., Broniewski, J., Westra, E., Bondy-Denomy, J., and Bassler, B.L. (2017). Quorum sensing controls the *Pseudomonas aeruginosa* CRISPR-Cas adaptive immune system. Proc. Natl. Acad. Sci. 114, 131–135. 10.1073/pnas.1617415113.

32. Li, C., Wen, A., Shen, B., Lu, J., Huang, Y., and Chang, Y. (2011). FastCloning: a highly simplified, purification-free, sequence-and ligation-independent PCR cloning method. BMC Biotechnol. 11, 92. 10.1186/1472-6750-11-92.

33. Lazarus, J.E., Warr, A.R., Kuehl, C.J., Giorgio, R.T., Davis, B.M., and Waldor, M.K. (2019). A new suite of allelic-exchange vectors for the scarless modification of proteobacterial genomes. Appl. Environ. Microbiol. 85, e00990–19. 10.1128/AEM.00990-19.

34. Ferrières, L., Hémery, G., Nham, T., Guérout, A.-M., Mazel, D., Beloin, C., and Ghigo, J.-M. (2010). Silent mischief: Bacteriophage Mu insertions contaminate products of <Escherichia coli<i> random mutagenesis performed using suicidal transposon delivery plasmids mobilized by broad-host-range RP4 conjugative machinery. J. Bacteriol. 192, 6418–6427. 10.1128/JB.00621-10.

35. Chimalapati, S., De Souza Santos, M., Servage, K., De Nisco, N.J., Dalia, A.B., and Orth, K. (2018). Natural Transformation in *Vibrio parahaemolyticus*: a Rapid Method To Create Genetic Deletions. J. Bacteriol. 200. 10.1128/JB.00032-18.

36. Hershey, A.D., Kalmanson, G., and Bronfenbrenner, J. (1943). Quantitative methods in the study of the phage-antiphage reaction. J. Immunol. 46, 267–279. 10.4049/jimmunol.46.5.267.

37. Sargen, M.R., Antine, S.P., Grabe, G.J., Antonellis, G., Ragucci, A.E., Li, Y., Kranzusch, P.J., and Helaine, S. (2026). A prophage-encoded abortive infection protein preserves host and prophage spread. Nature. 10.1038/s41586-025-10070-6.

38. Chiang, S.L., and Rubin, E.J. (2002). Construction of a mariner-based transposon for epitope-tagging and genomic targeting. Gene 296, 179–185. 10.1016/S0378-1119(02)00856-9.

39. Truong, T.T., Vettiger, A., and Bernhardt, T.G. (2020). Cell division is antagonized by the activity of peptidoglycan endopeptidases that promote cell elongation. Mol. Microbiol. 114, 966–978. 10.1111/mmi.14587.

40. Lan, S.-F., Huang, C.-H., Chang, C.-H., Liao, W.-C., Lin, I.-H., Jian, W.-N., Wu, Y.-G., Chen, S.-Y., and Wong, H.-C. (2009). Characterization of a new plasmid-like prophage in a pandemic *Vibrio parahaemolyticus* O3:K6 strain. Appl. Environ. Microbiol. 75, 2659–2667. 10.1128/AEM.02483-08.

41. Makino, K., Oshima, K., Kurokawa, K., Yokoyama, K., Uda, T., Tagomori, K., Iijima, Y., Najima, M., Nakano, M., Yamashita, A., et al. (2003). Genome sequence of *Vibrio parahaemolyticus: a pathogenic* mechanism distinct from that of *V. cholerae*. The Lancet 361, 743–749. 10.1016/S0140-6736(03)12659-1.

42. Simon, R., Priefer, U., and Pühler, A. (1983). A broad host range mobilization system for in vivo genetic engineering: Transposon mutagenesis in gram negative bacteria. Bio/Technology 1, 784–791. 10.1038/nbt1183-784.

43. Sargen, M.R., and Helaine, S. (2024). A prophage competition element protects Salmonella from lysis. Cell Host Microbe 32, 2063–2079.e8. 10.1016/j.chom.2024.10.012.

44. Lan, S.-F., Huang, C.-H., Chang, C.-H., Liao, W.-C., Lin, I.-H., Jian, W.-N., Wu, Y.-G., Chen, S.-Y., and Wong, H. (2009). Characterization of a new plasmid-like prophage in a pandemic *Vibrio parahaemolyticus* O3:K6 strain. Appl. Environ. Microbiol. 75, 2659–2667. 10.1128/AEM.02483-08.

